# Replication stress induced by CCNE1 overexpression creates a dependency on XRCC2 at the replication fork

**DOI:** 10.1101/325563

**Authors:** Kai Doberstein, Alison Karst, Paul T Kroeger, Ronny Drapkin

## Abstract

Across multiple cancer types, genome instability has been linked to aberrant over-expression of *CCNE1* due to premature cell cycle entry and replication stress. Using a gain-of-function screen, we found that *XRCC2* cooperates with *CCNE1* in the neoplastic transformation of *TP53* mutant cells. A pan-cancer analysis of TCGA data revealed a striking correlation between *CCNE1* and *XRCC2* expression and knockdown of XRCC2 in Cyclin E1 overexpressing cell lines is synthetic lethal. Immunopurification of XRCC2 showed that it interacts with the Minichromosome Maintenance Complex Component 7 (MCM7) protein. This interaction appears to be critical for protecting replication forks as knockdown of XRCC2 leads to a strong increase in MCM7 ubiquitination with concomitant decrease in MCM7 protein levels, and reduced replication fork speed. Importantly, Overexpression of MCM7 rescues the effect of XRCC2 knockdown. Our data describe a new dependency of Cyclin E1 overexpressing tumors on factors that stabilize the replication fork.

## INTRODUCTION

Ovarian cancer is the fifth leading cause of cancer mortality among women in developed countries. The 5-year survival rate is low and treatment options have not significantly improved in nearly two decades (Ferlay et al., 2015). Heterogeneity is a hallmark of this disease with high-grade serous ovarian carcinoma (HGSOC) being the most common subtype. Despite the lack of curative therapies, our understanding of HGSOC pathogenesis has greatly improved in the last 10 years. It is now generally accepted that a majority of HGSOCs arise from the fallopian tube (FT) secretory epithelial cell (Cancer Genome Atlas Research, 2011; Gorringe et al., 2010; Kroeger and Drapkin, 2017; Lee et al., 2007). Genomically, HGSOC exhibits a complex terrain with copy number alterations (CNA) rather than recurrent somatic mutations as the dominant aberration (Ahmed et al., 2010; Patch et al., 2015). The complex genomic landscape of HGSOC reflects defects in genome integrity control and DNA repair pathways with approximately 50% of cases manifesting dysfunction in the BRCA pathway and homologous recombination (HR). HR-deficient tumors are generally sensitive to first-line platinum therapy and PARP inhibition, which targets the compensatory DNA repair pathway (Fong et al., 2009; Scott et al., 2015). The remaining 50% of cases are equally complex but appear to have intact DNA repair. A significant fraction of these HR-proficient tumors express high levels of Cyclin E1, either through gene amplification or elevated expression. Interestingly, *BRCA1/BRCA2* mutations are mutually exclusive with *CCNE1* amplification and the knock-down of *BRCA1* in *CCNE1* amplified cells is synthetic lethal, indicating a dependency on the HR pathway in *CCNE1* amplified tumors (Etemadmoghadam et al., 2013). In contrast to HR-deficient patients, tumors harboring *CCNE1* amplifications are less likely to respond to primary platinum or PARP inhibitor treatment and have a reduced overall survival (Etemadmoghadam et al., 2009; Etemadmoghadam et al., 2013; Scott et al., 2015).

Our laboratory recently demonstrated that Cyclin E1 dysregulation manifests early during serous tumorigenesis with overexpression occurring in early FT precursors known as ‘p53 signatures’ and in serous tubal intraepithelial carcinomas (STIC) (Karst et al., 2014). Modeling these findings *in vitro* using immortalized FT secretory epithelial cells (FTSEC) revealed that the combined overexpression of mutant p53 and Cyclin E1 led to increased proliferation, loss of contact inhibition, clonogenic growth, and increased DNA damage (Karst et al., 2014). Interestingly, we observed only modest anchorage independent growth under these conditions. However, the Cyclin E1-overexpressing FTSECs demonstrated an upregulation of DNA repair factors that are essential for maintaining the integrity of the DNA replication fork, including BRCA1, FANCD2, BLM, CDC25, and XRCC2 (Karst et al., 2014). This response is consistent with premature entry into S phase of the cell cycle and the replicative stress imposed by Cyclin E1 overexpression (Chellappan et al., 1991; DeGregori et al., 1995; Hinds et al., 1992). Specifically, the aberrant expression of Cyclin E1 leads to a shortening of the G1-phase of the cell cycle, denying the cell adequate time to prepare resources that are needed for proper DNA replication during S-phase, such as sufficient nucleosides supply and proper origin licensing (Bester et al., 2011; Ekholm-Reed et al., 2004; Geng et al., 2003). Additionally, It has been shown recently that the activation of Cyclin E1 activates the firing of ectopic origins which are in conflict with transcription, resulting in replication stress and fork collapse (Macheret and Halazonetis, 2018). These effects notwithstanding, Cyclin E1 overexpression was not sufficient to transform FTSEC in the presence of mutant p53 alone.

In this report, we deployed a gain-of-function screen to identify genes that could fully transform FTSECs in the presence of mutant p53 and Cyclin E1 overexpression. We identified the RAD51 paralog XRCC2 as a gene that cooperates with CCNE1 to transform cells. We showed that XRCC2 plays a critical role in cells overexpressing Cyclin E1 by maintaining fork stability and enabling replication restart. XRCC2 achieves these effects by interacting and protecting the DNA replication licensing factor MCM7 at the stalled replication fork.

## RESULTS

### XRCC2 expression is strongly associated with Cyclin E1 expression and CNA

To better understand how Cyclin E1 contributes to the transformation of normal FTSECs, we utilized a previously described FT cell line model (Karst and Drapkin, 2012; Karst et al., 2014; Karst et al., 2011). The FT cell line was generated from human fallopian tube epithelium that was dissociated, immortalized with hTERT to delay senescence, and transduced with TP53^R175H^, one of the most common conformational *TP53* mutants identified in HGSOC and STIC lesions (Karst et al., 2011). The resulting untransformed, but p53-compromised FTSEC cell line model was named FT282.

To study the effects of constitutive Cyclin E1 expression in these cells, we transduced the FT282 cells with CCNE1 (FT282-CE) or a control vector (FT282-V) (Figure 1A) (Karst et al., 2014). Constitutive overexpression of Cyclin E1 imparted malignant characteristics on the cells, including increased colony formation, loss of contact inhibition, and more rapid proliferation. Additionally, we observed an increase in DNA damage as measured by γH2AX staining and an increase in comet assay tail length (Karst et al., 2011). Nevertheless, when cultured under anchorage independent conditions or soft agar, the Cyclin E1-expressing cells did not form robust colonies, suggesting that the combination of hTERT, mutant p53, and Cyclin E1 are insufficient to fully transform these cells.

**Figure 1:**
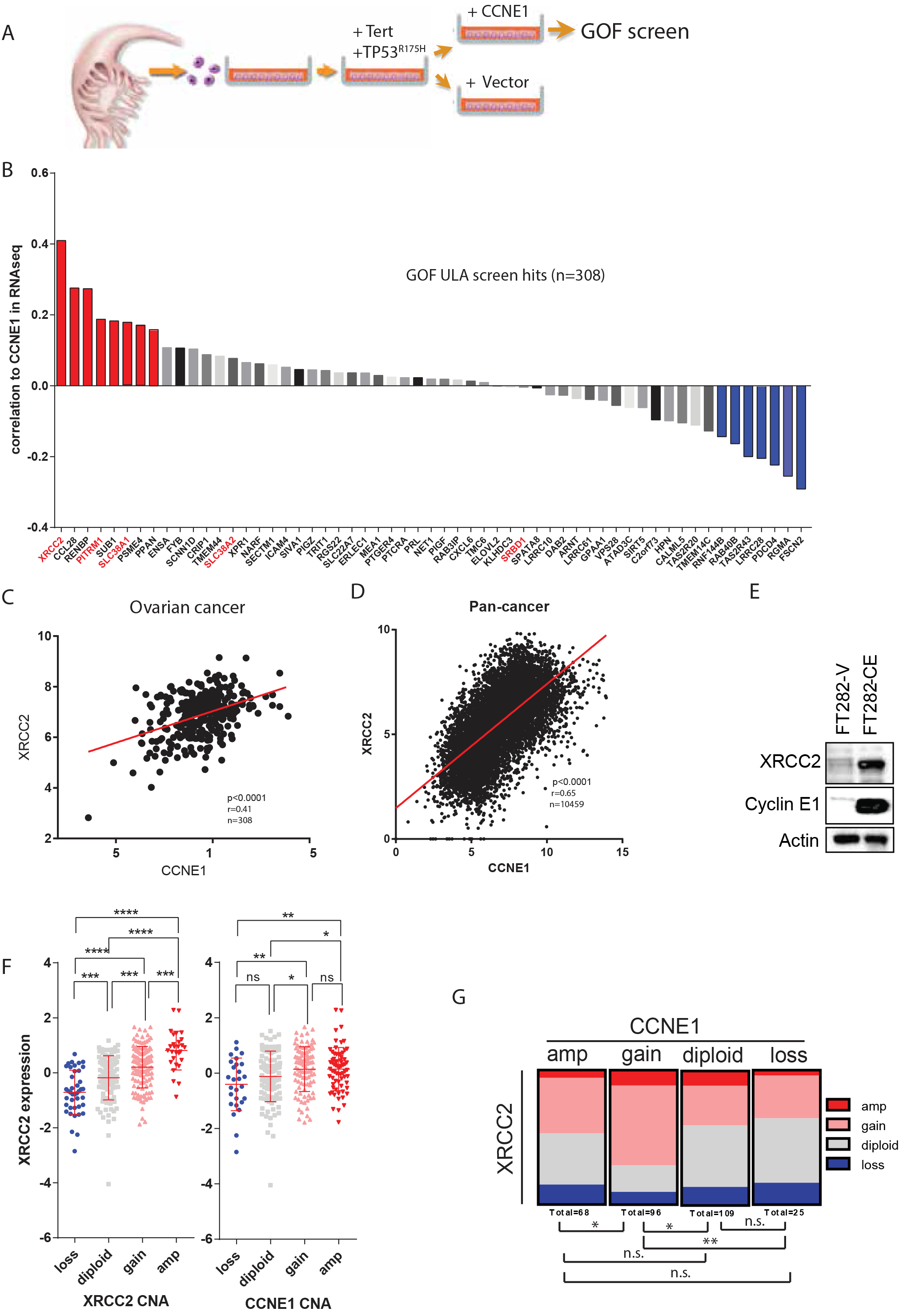
XRCC2 expression is strongly associated with Cyclin E1 expression and CNA. A) Schematic depicting the sequence of alterations used to generate primary human FTSECs (Karst et al., 2014). Cells were first immortalized with TERT and mutant TP53^R175H^, followed by the overexpression of *CCNE1* (FT282-CE) or a control vector (FT282-V). The Cyclin E1 overexpressing cells were then used for the anchorage independence gain of function (GOF) screen. B) Pearson correlation of the TCGA ovarian cancer cohort comparing RNAseq expression of GOF hits against *CCNE1* expression (n=308). Red columns indicate a significant (p<0.05) positive correlation, whereas blue columns indicate a significant (p<0.05) negative correlation. Fonts of gene names overlapping with the synthetic lethality screen are marked red. C) Scattered blot of the RNAseq expression of *XRCC2* against *CCNE1* expression in the TCGA ovarian cancer cohort (Pearson, r=0.41,p<0.0001) and D) Pan-cancer cohort (Pearson, r=0.65, p<0.0001), respectively. E) Western blot comparing differential protein expression of Cyclin E1 and XRCC2 in normal and Cyclin E1 overexpressing FTSEC cells. F) Relationship between *XRCC2* RNAseq expression and *XRCC2* CNA or *CCNE1* copy number alterations gathered from the TCGA ovarian cancer cohort. Indication of significance via p-values are described in material and methods. G) Relationship between *XRCC2* and *CCNE1* CNA extracted from the ovarian cancer TCGA data.

In order to identify genes that cooperate with Cyclin E1 to fully transform the FT282-CE cells, we performed a gain-of-function (GOF) screen in the FT282-CE cell line with a library of approximately 600 genes that are frequently amplified in ovarian cancer (Figure 1A) (Dunn et al., 2014). After transduction, we measured the ability of the cells to grow under anchorage independent conditions by analyzing colony size. We identified 57 genes whose overexpression led to anchorage independent growth of FT282-CE cells (Figure 1B, Figure S1A, B). Interestingly, five of those hits (*XRCC2, PITRM1, SLC38A1, SLC38A2* and *SRBD1*) overlapped with a recently performed synthetic lethality screen in *CCNE1* overexpressing cells, indicating that tumor cells likely create a dependency on those genes (Figure S1C, D) (Etemadmoghadam et al., 2013). We additionally analyzed the RNA expression of these 57 genes in The Cancer Genome Atlas (TCGA) for ovarian cancer and correlated them to *CCNE1* expression. This analysis revealed that *XRCC2* has the strongest correlation with *CCNE1* (r=0.41, Figure 1B, 1C, Figure S1E) and that this correlation is even stronger when analyzing the TCGA Pan-Cancer cohort (r=0.65,Figure 1D, Figure S1F).

XRCC2 belongs to the RAD51 paralog protein family that has been shown to be involved in HR and replication fork protection (Johnson et al., 1999; Somyajit et al., 2015). In agreement with our previously performed PCR array, we found that XRCC2 is also upregulated at the protein level in Cyclin E1 overexpressing fallopian tube cells (Figure 1E) (Karst et al., 2014). The strong correlation between *XRCC2* and *CCNE1* expression led us to examine whether the two genes are associated at the genomic level as well. We queried the TCGA ovarian cancer cohort for *XRCC2* RNAseq expression and its correlation to CNA. We analyzed CNA of *XRCC2* and *CCNE1* and found that there is a positive correlation between *XRCC2* RNA expression and increased CNA in both genes (Figure 1F). When comparing the CNA of *XRCC2* with *CCNE1* we found that most *XRCC2* gains and amplifications occur in samples with a gain of *CCNE1* (Figure 1G). These results, from two independent screens and the analysis of the TCGA data suggested a strong biological connection between *XRCC2* and *CCNE1* expression in HGSOC which we further investigated.

### XRCC2 knockdown is synthetic lethal in *CCNE1* overexpressing cells

In order to characterize the expression of XRCC2 and Cyclin E1 in HGSOC cells, we used a panel of cell lines that were previously analyzed for their CNA status of *CCNE1* by Western blot (Figure 2A) (Domcke et al., 2013). Additionally, we compared Cyclin E1 protein expression to the *CCNE1* copy number alterations and RNA expression from the Cancer Cell Line Encyclopedia (CCLE) that show a similar expression pattern compared to our Western blot analysis (Figure 2A). Interestingly, cell lines OVCAR4 and OVSAHO, which harbor a *CCNE1* copy number gain, express similar levels of *CCNE1* RNA as lines previously identified as harboring a *CCNE1* copy number amplification. Conversely, cell lines OVCAR8 and Kuramochi, which also harbor *CCNE1* copy number gain, express lower levels of *CCNE1* (Figure 2A), indicating a high variability of Cyclin E1 expression in cells harboring a *CCNE1* gain. To determine the effects of XRCC2 loss in these cells, we used siRNA to knock down XRCC2 (Figure 2B, Figure S2A, B). We found that knockdown of XRCC2 led to a strong reduction in cell survival in the cells that exhibited the highest levels of Cyclin E1 protein: OVCAR3, OVCAR4 and OVSAHO. Interestingly, when we plotted relative survival after XRCC2 knockdown against *CCNE1* copy number level, the survival measurement did not show a strong negative correlation with copy number level (Figure 2C). However, when we analyzed relative survival against *CCNE1* RNA or protein expression, we identified a significant negative correlation, indicating that the synthetic lethality of *XRCC2* knockdown is more dependent on overall Cyclin E1 protein expression than its *CCNE1* amplification status. These results are in agreement with the data from a synthetic lethality screen showing a dependency on *XRCC2* in Cyclin E1 overexpressing tumor cells (Etemadmoghadam et al., 2013). To analyze the dependency of Cyclin E1 overexpressing cells on XRCC2 in other cancer types, we investigated the dependency score of XRCC2 with the DepMap database (Cancer Dependency Map) that combines genome-wide loss-of-function screens (Achilles) with the molecular characterization of the CCLE and in which a negative score represents a stronger dependency of Cyclin E1 overexpressing cells on XRCC2 (Figure S2C)(Cowley et al., 2014; Tsherniak et al., 2017). We did not observe a significant correlation between XRCC2 dependency and overall Cyclin E1 RPPA or CNA levels using linear regression or pearson correlation (Figure S2D). However, when comparing the cells that expressed high versus low levels of Cyclin E1 protein (RPPA>1/O1) or *CCNE1* CNA levels (CNA>0.45/O0.45) we observed a significant difference in XRCC2 dependency (Figure S2E). In particular we observed a strong XRCC2 dependency in stomach cancer lines with high *CCNE1* CNA levels. A trend towards significance was also observed in endometrial, ovarian, pancreatic and prostate cancers (Figure S2F, G). Those data indicate that cell lines overexpressing Cyclin E1 are more dependent on XRCC2 expression.

**Figure 2:**
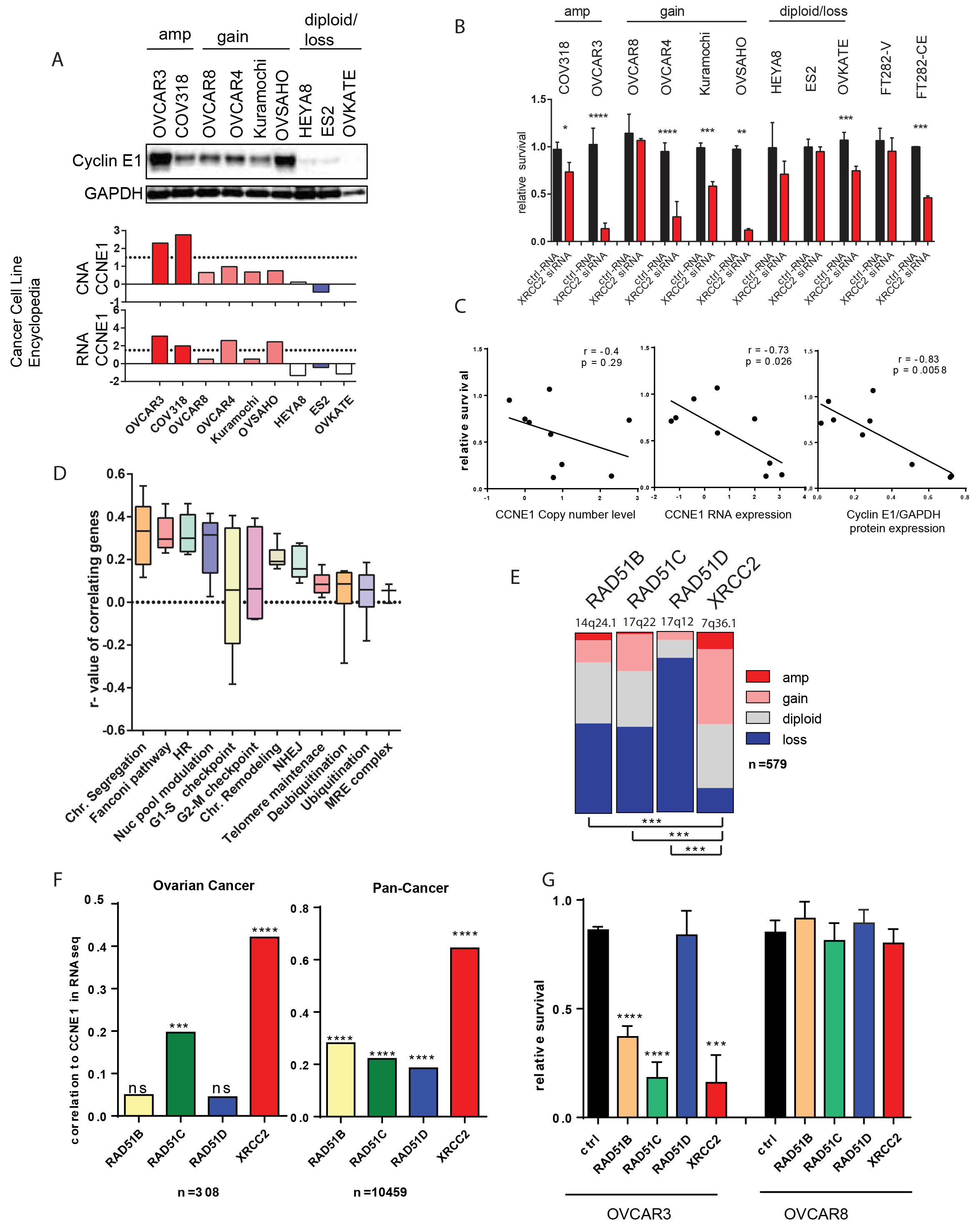
XRCC2 knockdown is synthetic lethal in *CCNE1* overexpressing cells. A) Upper row: Western blot analysis of Cyclin E1 in a cell line panel that was chosen using previously described Cyclin E1 levels (Domcke et al., 2013). Middle and lower row respectively: Relative *CCNE1* CNA and RNA levels from the Cancer Cell Line Encyclopedia (CCLE) in the cell lines analyzed. B) Survival analysis after transfection with *XRCC2* siRNA in cell lines after 72 hours relative to control siRNA. Strongest effect was observed in OVCAR3, OVCAR4 and OVSAHO. C) Relative survival from C was plotted against *CCNE1* copy number level (r=-0.4, p=0.29), RNA expression (r=-0.73, p=0.026) shown in B, or cyclin E1 protein expression (r=-0.83, p=0.0058) analyzed by densitometry (Cyclin E1/GAPDH) from Western blots shown in A. D) Genes of respective pathways were analyzed for RNAseq expression and their correlation to CCNE1 expression based on the ovarian cancer TCGA cohort. E) Copy number distribution of *RAD51B, RAD51C, RAD51D* and *XRCC2* in the ovarian cancer TCGA cohort. F) Correlation of the RNAseq expression of *CCNE1* against *RAD51B, RAD51C, RAD51D* and *XRCC2* in the TCGA ovarian cancer and pan-cancer cohort. G) Survival analysis after transfection with *RAD51B, RAD51C, RAD51D* or *XRCC2* siRNA in OVCAR3 and OVCAR8 cell lines after 72 hours relative to control siRNA.

To identify the cell cycle pathways that are commonly upregulated together with *CCNE1* overexpression, we performed a gene ontology analysis of cell cycle related pathways in the TCGA ovarian cancer dataset. We found that the genes involved in chromosome segregation, Fanconi anemia and homozygous recombination are strongly upregulated with *CCNE1* overexpression (Figure 2D). Since XRCC2 interacts with the other RAD51 paralogs, RAD51B/C/D, we also analyzed the paralogs because they are known to have roles in HR and replication fork stability (Compton et al., 2010). When analyzing the ovarian cancer and Pan-cancer TCGA RNAseq dataset, *XRCC2* expression correlated strongest with *CCNE1* expression, while *CCNE1* expression showed no correlation in the ovarian cancer dataset to *RAD51B* or *RAD51D* (Figure 2F). Perhaps more importantly, the CNA profile shows that among the paralogs, *XRCC2* has the most amplifications or gains and the least amount of losses in ovarian cancer (Figure 2E). Surprisingly, 84% of HGSCs had homozygous or heterozygous loss of *RAD51D* and no tumor harbored *RAD51D* amplifications (Figure 2E). Interestingly, we observed no effect of RAD51D knockdown in either cell line, suggesting that *RAD51D* might have a different role outside the traditional roles of the other *RAD51* paralogs in HGSOC (Figure 2G, Figure S3A). Those data indicate that cancer cells expressing high levels of Cyclin E1 develop a dependency on XRCC2 expression and, in this setting, loss of XRCC2 leads to cell death.

### XRCC2 knockdown leads to a decrease in replication speed in the context of Cyclin E1

One of the hallmarks of Cyclin E1 overexpression is the induction of genomic instability through DNA replication stress (Bester et al., 2011; Jones et al., 2013). In the setting of Cyclin E1 overexpression, it is thought that replication stress is induced by shortening the length of the G1 phase of the cell cycle. Contraction of G1 leads to an early, unscheduled entry into S-phase in unprepared cells. Since these cells enter S-phase prematurely, they have insufficient time to amass all the resources necessary for normal replication, such as providing enough nucleotides or loading of the replication origins (Ekholm-Reed et al., 2004; Geng et al., 2003; Macheret and Halazonetis, 2018). Those alterations subsequently lead to an increase in replication stress at the fork (Neelsen et al., 2013a).

Previous research has implicated the RAD51 paralogs in stabilizing the replication fork (Somyajit et al., 2015). Therefore, we speculated that XRCC2 might be an important factor for fork stabilization in *CCNE1*-amplified cells. To analyze the effects of XRCC2 during ongoing replication, we performed a fiber length assay 48h after XRCC2 siRNA transfection to measure the speed of the ongoing replication and replication termination (Figure 3A). Our results show that knockdown of XRCC2 in *CCNE1*-amplified (OVCAR3) versus non-amplified (OVCAR8) cells leads to a strong reduction in replication speed, as measured by fiber length per unit time (Figure 3B, 3C). However, this effect was not accompanied by an increase in stalled forks (Figure S3B). Consistent with our OVCAR3 data, when we analyzed the FT282 cell line overexpressing Cyclin E1, we found a much stronger reduction in replication speed upon XRCC2 knockdown (Figure 3D). To be able to replicate the whole genome in a given amount of time, cells can compensate for the reduction in replication speed by firing extra origins, known as ‘dormant origins’ (Fragkos et al., 2015). Therefore, we analyzed the changes in origin distance after XRCC2 siRNA knockdown, and found a significantly shorter interval between origins (Figure 3E). Interestingly, the origin distance of the FT282 overexpressing Cyclin E1 was already significantly shortened compared to the control cells, suggesting *CCNE1* amplification causes the activation of dormant origins, even before knockdown of XRCC2. All these data indicate that XRCC2 plays an important role in maintaining normal replication rates in Cyclin E1 overexpressing cells.

**Figure 3:**
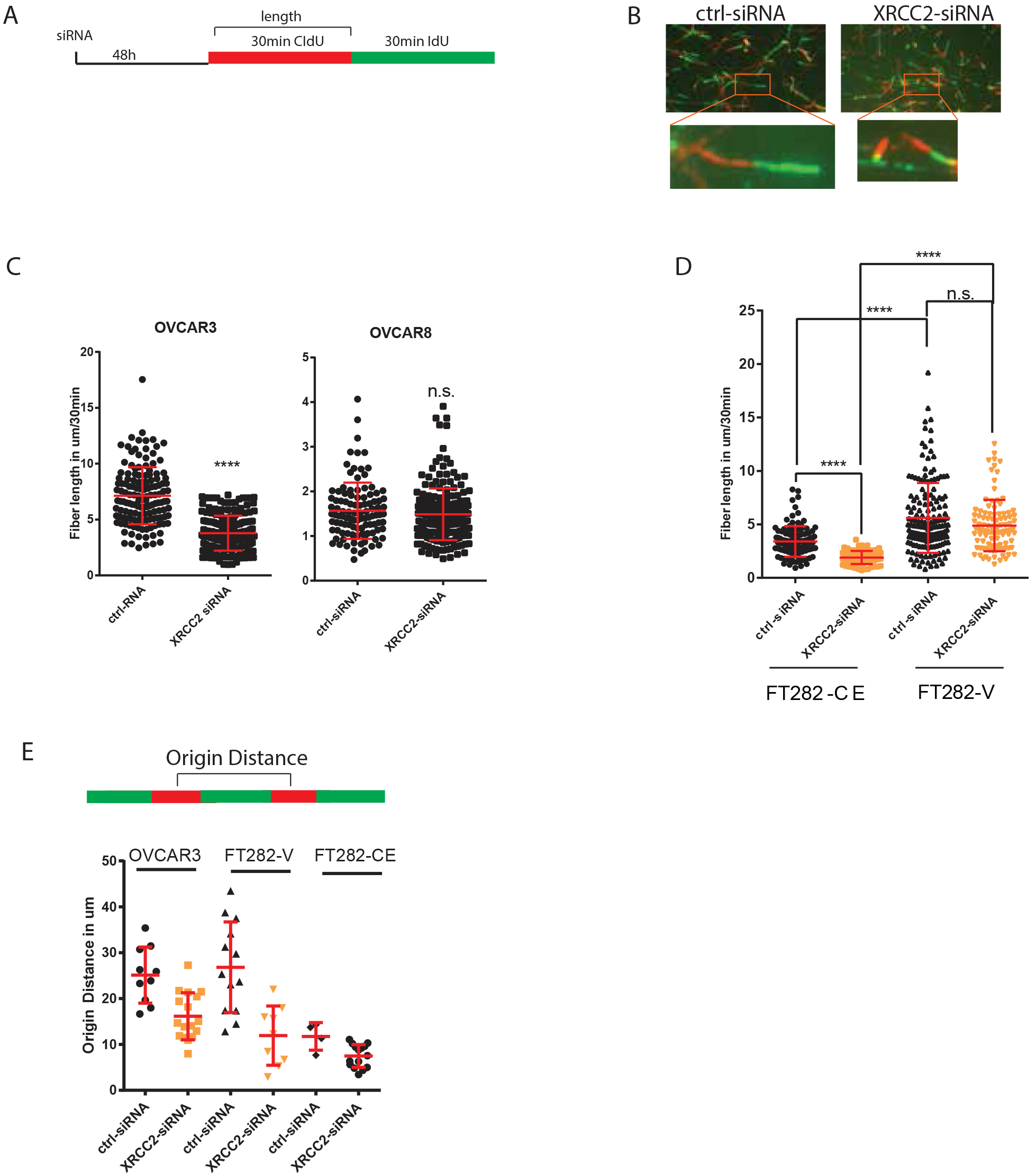
XRCC2 knockdown leads to a decrease in replication speed in the context of Cyclin E1 overexpression. A) An illustration of the experimental design of the fiber length assay. 48 hours after siRNA transfection cells were treated for 30 minutes with CIdU followed by 30 minutes of IdU incubation. CIdU length was measured in the following experiments. B) Representative fiber length analysis 48 hours after cells treated with control or *XRCC2* siRNA. C) Fiber length analysis of OVCAR3 and OVCAR8 cells after transfection with control siRNA or siRNAs against *XRCC2*. D) Fiber length measurements 48 hours after transfection of *XRCC2* and control siRNA in FT282-V and FT282-CE cells. E) Bar illustrates the measurement of the distance between origins, therefore the center of two neighboring CIdU labels were measured. The origin distance was determined following *XRCC2* or control siRNA transfection in OVCAR3, FT282-V and FT282-CE

### XRCC2 is important for replication fork recovery

The effect of XRCC2 knockdown on replication fork speed prompted us to analyze the role of XRCC2 at the replication fork. We tested which type of DNA stress cooperates synergistically with XRCC2 knockdown to further reduce the ability of the replication fork to recover. After XRCC2 siRNA treatment and incubation in CIdU for 30 minutes, we treated FT282-CE cells with the following DNA damaging agents for two hours: 1) ultraviolet light (UV) to generate DNA adducts, 2) hydroxyurea (HU), to deplete nucleotides at the replication fork, 3) neocarzinostatin (NCS), 4) doxorubicin (DOX) or 5) the DNA cross-linking agent cisplatin (CDDP). Following DNA damage, we added IdU for 30 minutes to assess the ability of the fork to recover (Figure 4A). Interestingly, treatment with UV, NCS or DOX led to an increase in fork speed recovery in XRCC2 knockdown cells due to the firing of new origins (Figure 4B, 4C). Only the treatment with HU led to a more dramatic shortening in fiber length, suggesting that the effects of XRCC2 are due to its role in replication fork recovery. We found a similar effect when cells were treated with HU for a longer time periods (Figure 4D).

**Figure 4:**
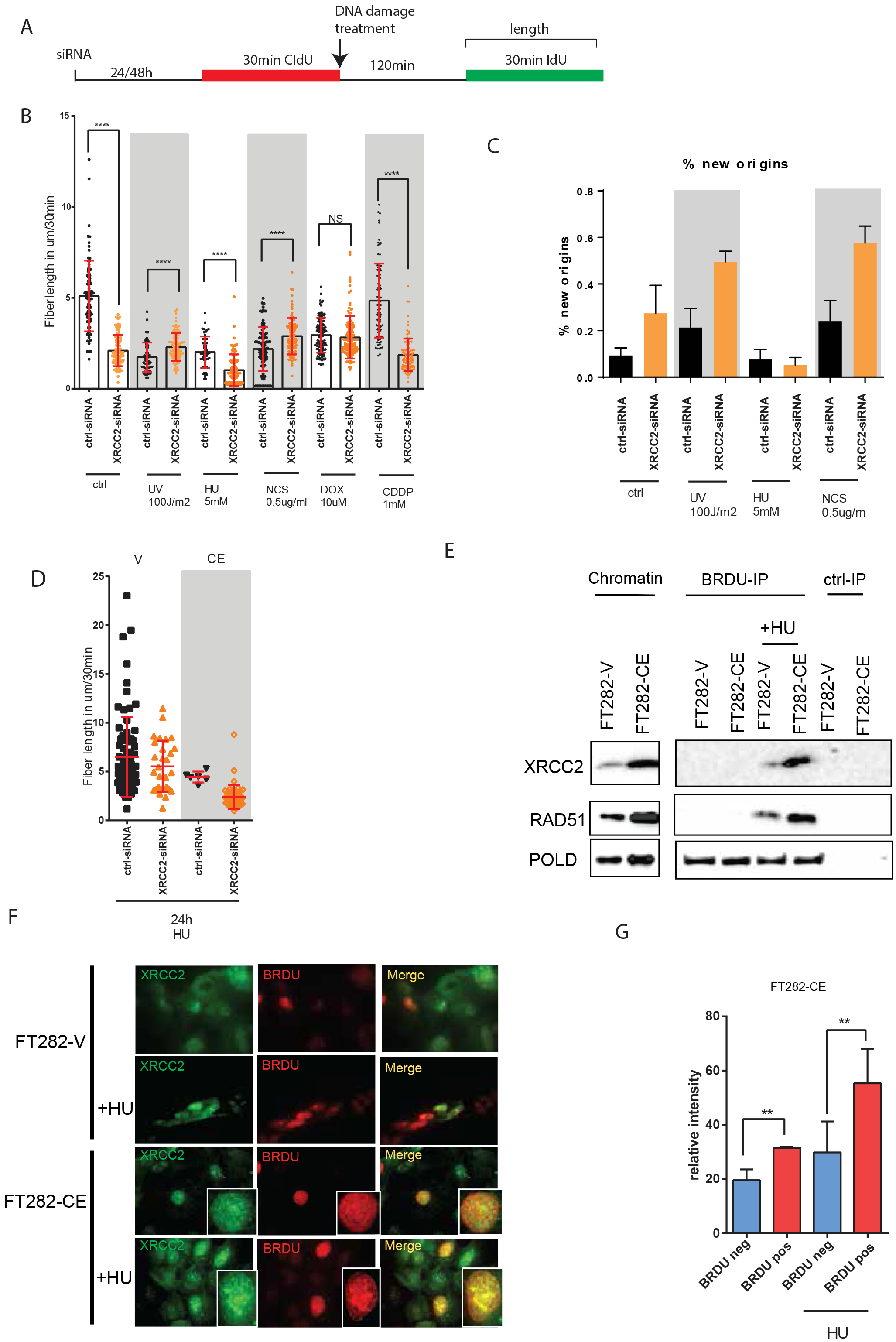
XRCC2 is important for replication fork recovery. A) Experimental schematic for DNA replication restart visualization including treatment with various DNA replication stressors. IdU fiber length was measured. B) IdU fiber length was measured as after treatment with Ultraviolet light (UV), Hydroxyurea (HU), Neocarzinostatin (NCS), Doxorubicin (Dox) or cisplatin (CDDP). C) The relative number of new origins were measured for control conditions UV, HU, NCS and Dox treated cells. D) IdU fiber length measured after 24 hours of treatment with HU in FT282-V and FT282-CE cells. E) FT282-V and FT282-CE cells were labeled with BrdU and treated with either HU for24 hours or directly treated with BrdU. Cells were cross-linked and chromatin fractionwas isolated and used for immunoprecipitation. Immunoprecipitation was performed against BrdU and analyzed by Western blot for XRCC2, RAD51 and POLD. F) Representative images of FT282-V and FT282-CE cells treated with HU and analyzedfor XRCC2 expression (green) and replicating DNA (BrdU (re D)). G) Quantitative analysis of XRCC2 florescence intensity in replicating cells (BrdU positivE) and non-replicating (BrdU negativ>E) FT282-V and FT282-CE cells.

Prior studies have shown that HU treatment leads to the recruitment of XRCC2 to the replication fork (Somyajit et al., 2015). Therefore, we used immunoprecipitation of BrdU labeled fork DNA to analyze the interaction of XRCC2 with the replication fork in FT282 cells. Treatment with HU for 24h led to increased binding of XRCC2 at the replication fork. Interestingly, the FT282-CE overexpressing Cyclin E1 cells recruited more XRCC2 to the replication fork (Figure 4E). We confirmed this observation with an immunofluorescence experiment that showed the co-localization of BrdU and XRCC2 after 24h HU treatment in FT282-CE vs FT282-V cells (Figure 4F). Additionally, we found that BrdU-positive cells expressed higher XRCC2 levels after HU treatment (Figure 4G). These data indicate that XRCC2 recruitment to the replication fork is important in cells that overexpress Cyclin E1, and that it enables these cells to recover after prolonged replication stress.

### XRCC2 interacts with MCM7 during DNA stress

To better understand how XRCC2 might contribute to the recovery of the replication fork in the context of *CCNE1* amplification, we sought to identify the factors that might interact with XRCC2 at the replication fork. After transiently transfecting GFP-tagged XRCC2 into OVCAR3 cells, we immunoprecipitated XRCC2 from the chromatin fraction with high-affinity GFP-Trap. GFP-tagged XRCC2 has been previously shown to be functional (O’Regan et al., 2001). Uncoupled magnetic agarose beads served as the negative control. The immunoprecipitated proteins were identified by mass spectrometry (Figure S4A). Interestingly, one of the XRCC2-interacting proteins was minichromosome maintenance complex component 7 (MCM7) (Figure 5A), a member of the replication machinery and a necessary component for the assembly of the MCM2-7 helicase (Labib et al., 2000). To confirm this interaction, we immunoprecipitated endogenous MCM7 and XRCC2 in FT282-CE and OVCAR3 cells and could thereby show that XRCC2 binds MCM7 in both cell lines (Figure 5B). To further validate the interaction between XRCC2 and MCM7, we used a proximity ligation assay (PLA) and measured the interactions by counting foci per cell. Using this method we counted significantly more interactions per nucleus in the FT282 cells overexpressing Cyclin E1 than in the control cells (Figure 5C), suggesting increased interactions in cells that overexpress Cyclin E1. To demonstrate that MCM7 and XRCC2 directly interact, we incubated recombinant MCM7 (rMCM7) with either GST-XRCC2 or GST alone. We observed that only GST-XRCC2 was able to directly interact with rMCM7 (Figure 5D). Interestingly, analysis of the TCGA ovarian cancer RNAseq expression data revealed that *XRCC2* had the highest correlation to *MCM7* (r=0.56) compared to other interacting proteins found in our mass spectrometry analysis (Figure 5E, Figure S4A-C). In addition to the TCGA ovarian cancer cohort, we also found a strong direct correlation between *MCM7* and *XRCC2* RNAseq expression in the pan-cancer cohorts (Figure 5E, Figure S4D).

**Figure 5:**
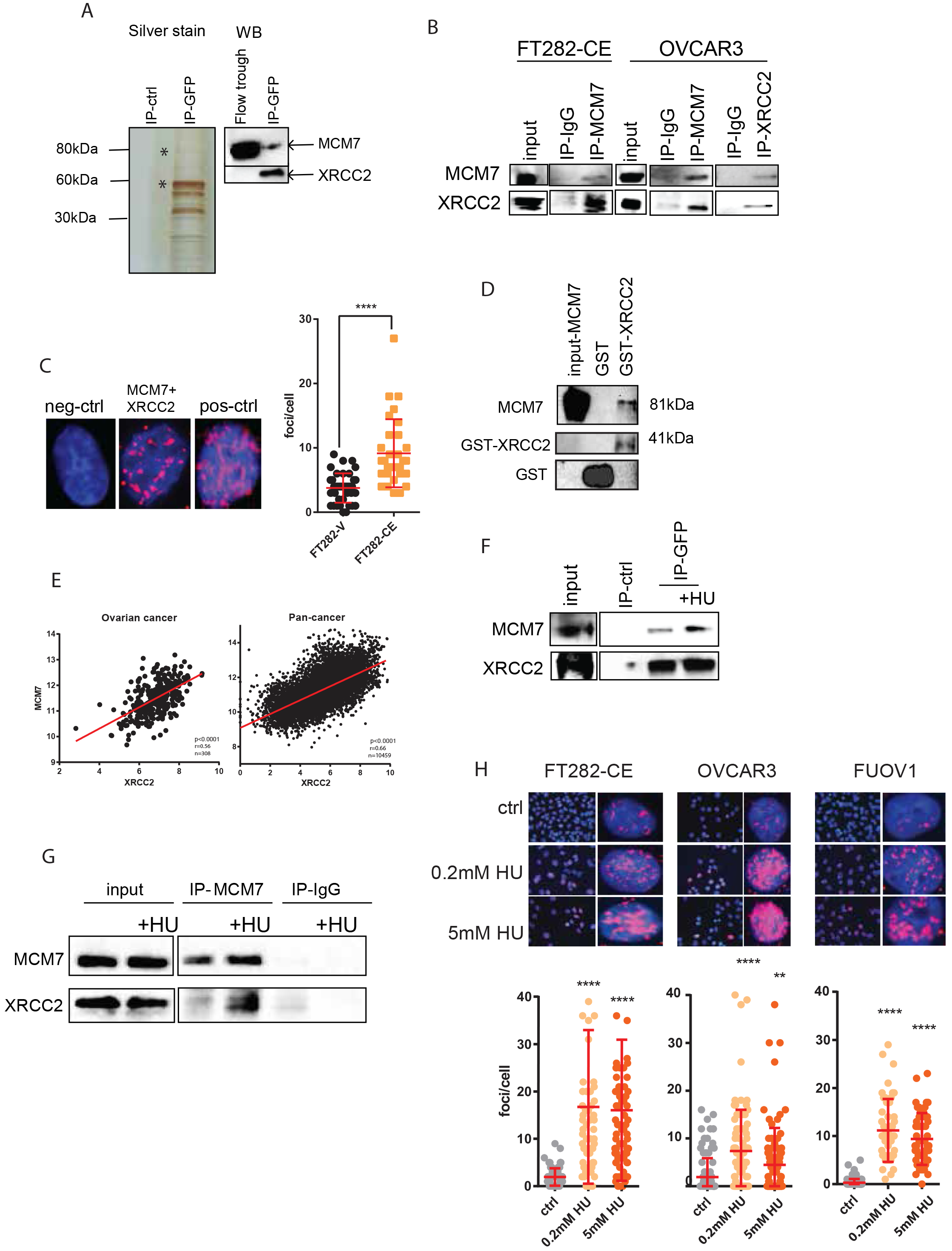
XRCC2 interacts with MCM7 during DNA stress. A) Silver stain of an anti-GFP pull-down of the nuclear fraction of OVCAR3 cells transiently transfected with XRCC2-GFP. Validation of pull-down was demonstrated by Western blot of MCM7 and XRCC2. B) The chromatin fraction of XRCC2 and MCM7 were immunoprecipitated from FT282-CE and OVCAR3 cells and blotted against MCM7 and XRCC2. C) Proximity ligation assay (PLA) between MCM7 and XRCC2 in FT282-V and FT282-CE cells. Cells were quantified by foci per cell. A representative picture of the experiment is shown together with a negative and positive control. D) GST pull-down of recombinant GST-XRCC2 incubated with recombinant MCM7 was analyzed by Western blot against XRCC2, MCM7 and GST. E) Scattered blot of the RNAseq expression of *MCM7* against XRCC2 expression in the TCGA ovarian cancer cohort and Pan-cancer cohort respectively. F) Western blot analysis of an anti-GFP pull-down from OVCAR3 cells transfected with XRCC2-GFP and treated with HU for 24 hours followed by blotting against MCM7 and XRCC2. G) Western blot analysis of an MCM7 pull-down from OVCAR3 cells with and without HU treatment for 24 hours, blotted for MCM7 and XRCC2. H) PLA between MCM7 and XRCC2 in FT282-CE, OVCAR3 and FUOV1 after treatment of 0.2mM and 5mM HU for 24 hours. Cells were analyzed by foci per cell.

In light of our results showing that XRCC2 plays a role at the replication fork during replicative stress, we tested whether the MCM7-XRCC2 interaction is influenced by treatment with HU using a co-immunoprecipitation assay. The resulting Co-IP showed an increased binding of MCM7 to XRCC2 in cells treated with HU for 24h. This was demonstrated by reciprocal IP’s using MCM7 and XRCC2-GFP (Figure 5F, 5G). Next, we asked whether the activation of dormant origins with low levels of HU could influence the interaction of MCM7 with XRCC2. To investigate this HU-induced interaction, we tested whether high and low doses of HU can increase the interaction of MCM7 and XRCC2 by PLA. We found a significant increase in foci after treatment with either concentration of HU, indicating that XRCC2 is likely involved in the firing of dormant origins under replication stress (Figure 5H). These data support the presence of a physical interaction between XRCC2 and MCM7 that is increased by replication stress.

### XRCC2 stabilizes and regulates the protein expression of MCM7

The strong correlation between *MCM7* and *XRCC2* in the ovarian cancer and pan-cancer TCGA dataset (Figure 5D), prompted us to further investigate the mutual dependency of both genes. When we compared the correlation of XRCC2 to the other MCM members we found a lower but significant correlation between XRCC2 and each MCM member (Figure 6A). When comparing it the other genes of the RAD51 paralogs family, XRCC2 expression clustered strongest with the expression of the MCM genes (Figure S4E). *CCNE1* and *MCM7* RNAseq expression also correlated positively in the TCGA ovarian cancer and the pan-cancer cohorts (r=0.29, r=0.63, respectively). *XRCC2* and *MCM7* are both located on chromosome 7 at positions 7q22.1 and 7q36.1, respectively. The distance between the genes is approximately 52 megabases (Mb). This is likely too far to be located within the same amplicon as studies have shown that the biggest amplicon on chromosome 7 has a size of approximately 4 Mb (Malek et al., 2011). Nevertheless, when analyzing the CNA of both genes, we found a high rate of co-occurrence of both amplifications and gains between the genes (Figure 6B). By analyzing the dependency of *XRCC2* and *MCM7* in the Achilles dataset, we could further show that cell lines that are dependent on *XRCC2* are also dependent on *MCM7* (Figure S5A-E)

**Figure 6:**
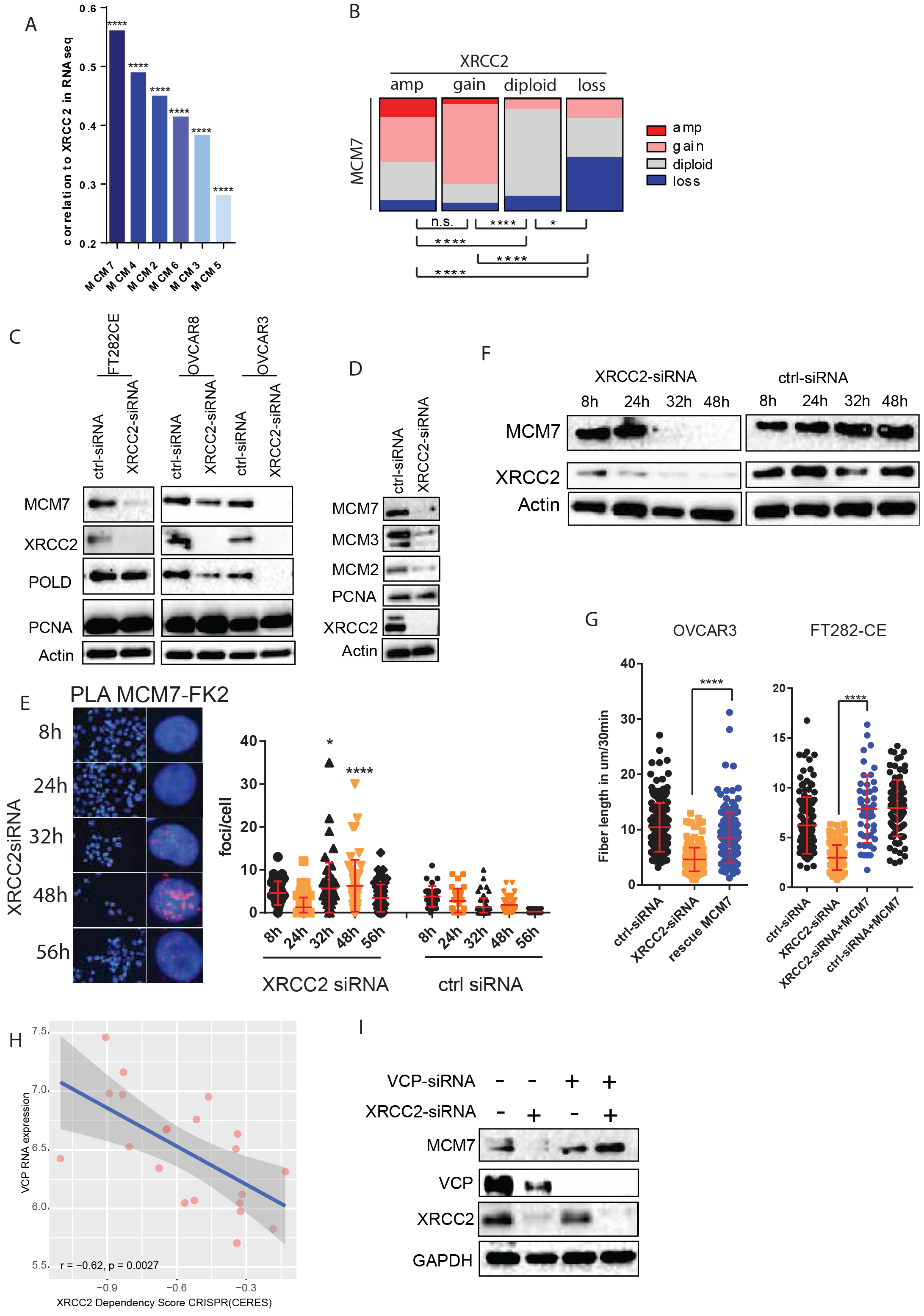
XRCC2 stabilizes and regulates the protein expression of MCM7. A) Histogram of correlation coefficients of the TCGA RNAseq expression of *MCM2-7* against *XRCC2* expression. B) Relationship between *XRCC2* and *MCM7* copy number alterations in the TCGA ovarian cancer cohort. C) Western blot analysis of MCM7, XRCC2, POLD and PCNA after siRNA knockdown of XRCC2 in FT282-CE, OVCAR8 and OVCAR3 cells. D) Western blot analysis of MCM7, MCM3, MCM2, PCNA, XRCC2 after siRNA knockdown in FT282-CE cells. E) Polyubiqitination of MCM7 was analyzed by PLA with antibodies against FK2 and MCM7 at various time points after XRCC2 siRNA knockdown in OVCAR3 cells. Cells were quantified by foci per nuclei. 8 hours before fixation, cells were treated with the proteasome inhibitor bortezomib. F) Western blot analysis of the time kinetic by blotting for MCM7 and XRCC2 at various time points after XRCC2 siRNA treatment in FT282-CE cells. G) Fiber length rescue analysis in OVCAR3 and FT282-CE measured 48 hours after transfection of XRCC2. 8 hours after siRNA transfection a MCM7 vectors was transfected. H) Scatter plot of *VCP* RNA expression against the *XRCC2* dependency score of the Achilles CRISPR(CERES) dataset in ovary adenocarcinoma cells (Pearson: r=-0.62, p=0.0027). I)Western blot analysis of MCM7, XRCC2, VCP and GAPDH after siRNA knockdown of XRCC2 and VCP.

Since both genes demonstrate a strong positive association, we asked whether there might be a direct dependency of MCM7 on XRCC2. Indeed, in multiple cell lines we found that the knockdown of XRCC2 led to a strong decrease in MCM7 expression (Figure 6C). The downregulation of one of the MCM proteins has been shown to lead to the destabilization of the other five members of the MCM hexamer complex (Moreno et al., 2014). We therefore investigated the stability of the MCM complex after the knockdown of XRCC2, and showed MCM2 and MCM3 were also downregulated together with MCM7 in FT282-CE cells (Figure 6D).

Previously, it has been shown that the protein INT6 is able to bind MCM7 to protect it from ubiquitination and thus from proteasomal degradation (Buchsbaum et al., 2007). Therefore, we used the PLA method to ask whether MCM7 is polyubiquitinated upon knockdown of XRCC2. We used antibodies that recognize the polyubiquitin chains (FK2) and MCM7 to detect polyubiqutinated MCM7. We performed a time course experiment and found that the ubiquitination of MCM7 is strongest 32 to 48 hours after siRNA knockdown of XRCC2 (Figure 6E). This result is in agreement with the strong reduction of MCM7 that we observe by western blot analysis 32 hour after the XRCC2 knockdown (Figure 6F). By using the proteasome inhibitors BZT and MG132, we were able to partially rescue the effects on replication fork speed in Cyclin E1 overexpressing cells (Figure S6A). Importantly, by using an MCM7 overexpressing cDNA, we were able to rescue the reduced speed phenotype at the replication fork that is generated by XRCC2 loss in *CCNE1* overexpression (Figure 6G). While the E3-ubiquitin ligase which ubiquitinates MCM7 at the chromatin has not been identified, it has been demonstrated that the process of replisome termination is facilitated through the VCP/p97 secretase and the K48-ubiquitination on MCM7 (Maric et al., 2014; Moreno et al., 2014). We therefore tested whether the effects of the XRCC2 knockdown is dependent on the expression of VCP. By analyzing the Achilles dataset we could show that ovarian cancer cells expressing higher levels of VCP are have a higher dependency on XRCC2 (Figure 6H, Figure S6B,C). By simultaneous downregulation of XRCC2 and VCP with RNAi, we were able to partially rescue the downregulation of MCM7 (Figure 6I), supporting the role of VCP in MCM7 stability. Similarly, INT6 which has been shown to protect MCM7 from ubiquitination shows a strong negative correlation with XRCC2 dependency (Figure S6D,E). Together, these data indicate that the interaction of MCM7 with XRCC2 has a protective role at the replication fork.

## DISCUSSION

Understanding the biology and vulnerabilities of *CCNE1*-amplified tumors is essential for the development of more effective therapies for this class of tumors. In this study, we used a gain-of-function screen to identify *XRCC2* as a gene that not only cooperates with CCNE1 to transform fallopian tube cells but is also synthetic lethal in *CCNE1*-amplified tumors. Among the various hits identified in the screen, *XRCC2* showed the highest positive correlation with *CCNE1*, not only in the ovarian-cancer cohort, but also in a pan-cancer analysis of the TCGA datasets. This indicates a connection in expression levels between both genes which is independent of cancer type. *XRCC2,* along with the other *RAD51* paralogs *RAD51B, RAD51C* and *RAD51D,* is best known for its role in the homologous recombination (HR) pathway of DNA repair (Johnson et al., 1999). XRCC2, together with the other paralogs, forms a complex (BCDX2) that functions downstream of RAD51 filament assembly by binding and remodeling RAD51 bound single stranded DNA filaments to be more flexible and thus enabling strand exchange (Miller et al., 2004; Taylor et al., 2015).

It has been shown that overexpression of Cyclin E1 leads to replication stress and DSBs (Neelsen et al., 2013a; Neelsen et al., 2013b). One proposed mechanism is that DSBs occur because of prolonged fork stalling and increased distances between replication forks. The reduced number of licensed origins lead to an increased origin distance that consequently creates fork collapse and DSBs (Letessier et al., 2011; Ozeri-Galai et al., 2011). Additionally, it has been recently demonstrated that the premature S phase entry, following the induction of CCNE1, activates replication origins within highly transcribed genes, resulting in a conflict between replication and transcription (Macheret and Halazonetis, 2018). Those CCNE1 activated origins are prone to DSB formation and chromosomal rearrangement. Interestingly, the paralogs complex, BCDX2, has been shown to preferably bind Y-shaped replication-like DNA as a ring like structure and to promote replication fork progression and restart after DNA damage (Compton et al., 2010; Henry-Mowatt et al., 2003; Yokoyama et al., 2004). Recently, it has been further demonstrated that the RAD51 paralogs are able to prevent DSB at the stalled replication fork and mediate continuous DNA synthesis (Somyajit et al., 2015). We therefore speculated that XRCC2 might play an important role in replication fork protection in cells that are constantly exposed to replication stress, including *CCNE1*-amplified cells. Indeed, we found that in the context of Cyclin E1 overexpression, XRCC2 knock-down leads to a marked replication slow down, origin distance shortening, and a reduced ability to recover from replication stress. Additional replication stress led to an increased recruitment of XRCC2 to the replication fork, indicating the importance of XRCC2 in fork protection and fork restart in Cyclin E1 overexpressing cells. Interestingly, we did not observe an increase in fork stalling upon XRCC2 knock down, but a strong reduction in restart speed, indicating that XRCC2 might be involved in the regulation of the replication fork speed by interacting with proteins involved with replication.

Importantly, we found that XRCC2 can bind MCM7. MCM7 is part of the DNA licensing hexamer MCM (MCM2-7) complex that functions as a helicase to unwind DNA at replication origins (Labib et al., 2000; Lei and Tye, 2001; You et al., 1999). We further found that the interaction of MCM7 and XRCC2 was augmented when we increased the stress at the replication fork by depleting nucleosides with HU. Consistent with our results, it has been shown that FANCD2 can also directly interact with the MCM complex upon replication stress and thereby prevent the accumulation of pathological replication structures (Lossaint et al., 2013). Similarly, RAD17 has been shown to interact with MCM7 after UV treatment to mediate ATR activation (Tsao et al., 2004). These data indicate that it is important for the MCM complex to sense and to respond to different types of DNA damage and replication stress.

Our data showed that XRCC2 stabilizes MCM7 during replication stress and that loss of XRCC2 leads also to a strong reduction of MCM7 expression. It might be possible that a stalled replication fork leads to conformational changes of the MCM complex that subsequently lead to its degradation if not protected. Consistent with this hypothesis, it was found that the protein INT6 stabilizes MCM7 from degradation (Buchsbaum et al., 2007). Importantly, it was further shown that MCM7 is the only MCM complex protein that is polyubiquitylated during replication and that the loss of one MCM protein leads to destabilization and degradation of the entire MCM complex (Moreno et al., 2014). Therefore, XRCC2 plays a crucial role in stabilizing the MCM7 protein and its MCM complex by binding and protecting it from degradation; an important function during replication stress.

Interestingly, Cyclin E1 has been associated with the loading of the MCM complex during the G1 phase to the chromatin (Chuang et al., 2009; Coverley et al., 2002). Nevertheless, when Cyclin E1 was strongly overexpressed, the G1-phase shortening led to an aberrant assembly of the pre-replication complex, resulting in a reduced number of properly licensed origins during S-phase (Ekholm-Reed et al., 2004; Geng et al., 2003). This subsequently led to an increased distance between replication forks, and by increasing the average replicon size, often resulted in unfinished chromosome replication before entry into mitosis (Letessier et al., 2011; Ozeri-Galai et al., 2011). As a consequence the replicated chromosomes undergo breakage, leading to a higher frequency of double-stranded DNA breaks and other karyotypic abnormalities.

Our data showed that the knockdown XRCC2 leads to a reduction in fork speed and origin distance. Since normal cells load MCM origins in excess, as a licensing backup to prepare for possible replicative stress, they are able to tolerate up to a 90% loss of MCM proteins and remain viable (Ge et al., 2007). In contrast, cells overexpressing Cyclin E1 cause a shortening of the G1 phase that ultimately leads to fewer origins licensing during the G1 phase. The resulting replication stress during S phase leads to the slowdown of the replication fork and the firing of more origins that are located closer to each other, consequently leading to a strong reduction of excess loaded origins for backup firing. A further reduction of MCM proteins, combined with the previously described replication stress, would therefore be difficult to tolerate for Cyclin E1 overexpressing cells. Furthermore, it has been shown that the suppression of MCM proteins makes cells hypersensitive to DNA replication stresses through increased frequency of chromosome breaks (Ibarra et al., 2008). Recently, it was demonstrated that Simvastatin can downregulate MCM7, which led to increased DNA damage and cell death (Liang et al., 2017). It would be interesting to combine MCM reducing drugs, like Simvastatin, with other DNA damaging agents, to therapeutically treat CCNE1-amplified HGSOC patients.

In summary, our experimental data have identified some of the molecular mechanisms whereby XRCC2 cooperates with CCNE1 to transform FT cells and initiate tumorigenesis.

## Acknowledgments

We thank members of the Drapkin lab for fruitful discussions and comments. We thank Roger Greenberg and Qinqin Jiang for their help to establish the fiber spread assay. We thank William Hahn and Gavin Dunn from the DFCI and the Broad Institute for the gain-of-function screen library and protocol. We would like to thank Matthew D. Weitzman, Rugang Zhang and Pat Morin for constructive criticism of the manuscript. This work was supported by the German Research Foundation DFG (K.D.), NIH P50-CA083636 (R.D.), the Dr. Miriam and Sheldon G. Adelson Medical Research Foundation (R.D.), the Honorable Tina Brozman Foundation for Ovarian Cancer Research (R.D.), The Basser Center for BRCA (R.D.), and the Claneil Foundation (R.D.).

## Author Contributions

Author contributions: K.D. and R.D developed the concept and design of the research; K.D., A.K. and P.T.K. performed the research; K.D. and R.D. analyzed the data; K.D. and R.D. wrote the paper, and all authors reviewed and edited the manuscript

## Supplementary Figure legends

**Figure S1.**
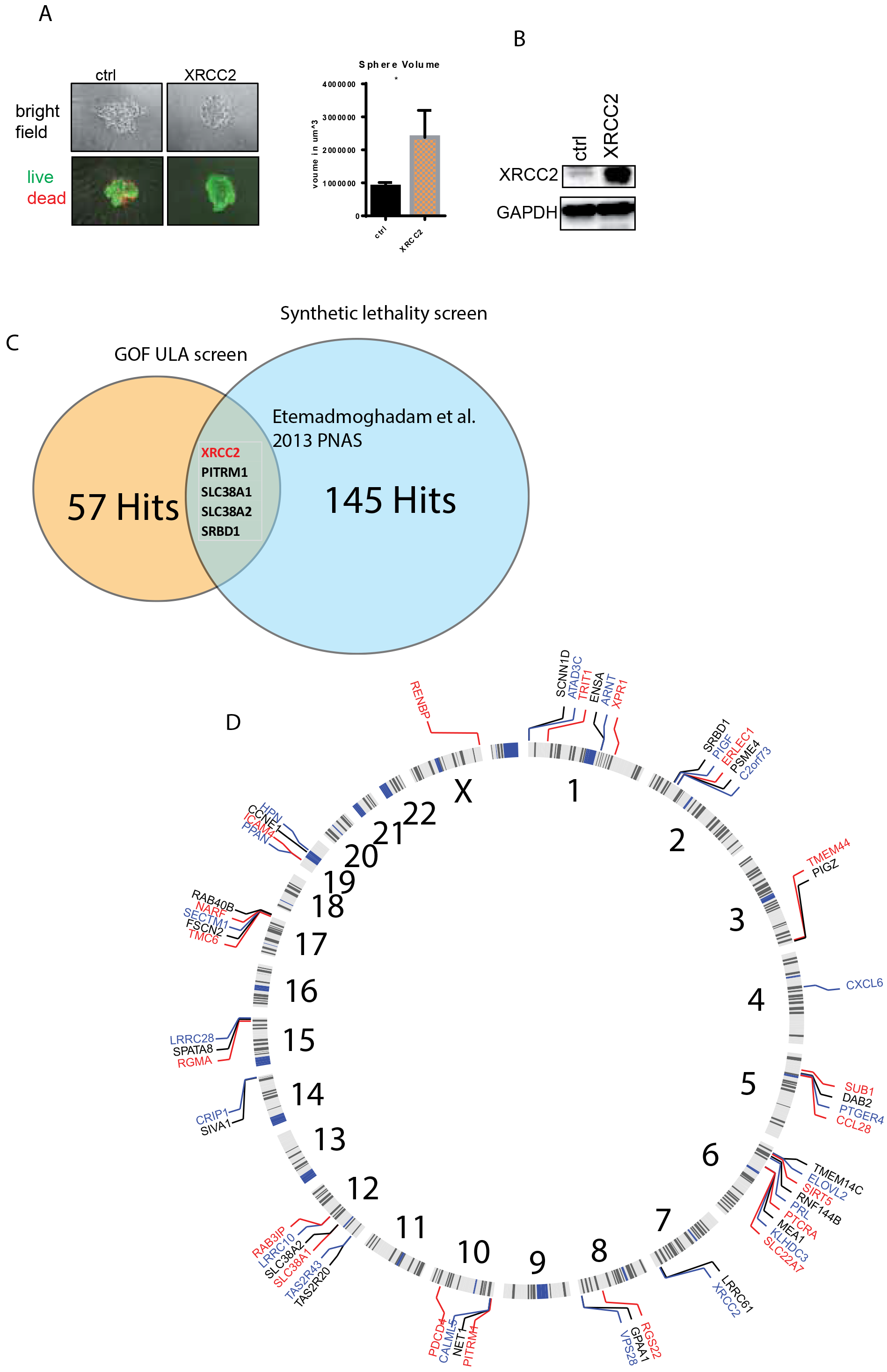
A) Left: Bright field image (upper panel) and live/dead staining (lower panel, green: living, red :dead) of sphere growth under ULA conditions of FT282-CE cells transfected with a control vector or with a vector expressing XRCC2. Right: Quantification sphere volume. B) Western blot analysis of FT282-CE cells transfected with a control vector or with a vector expressing XRCC2. C) Venn diagram of the overlapping genes of our GOF screen and collaborative synthetic lethality screen (Etemadmoghadam et al., 2013) D) Diagram shows the genomic location of the 57 identified genes. E) Correlation matrix of the RNAseq expression of the 57 identified genes based on the TCGA ovarian cancer dataset and hierarchical clustered. F) Scattered blot of the RNAseq expression of *XRCC2* against *CCNE1* expression in the TCGA Pan-cancer cohort separated for each individual cancer type. Depicted is pearson correlation.

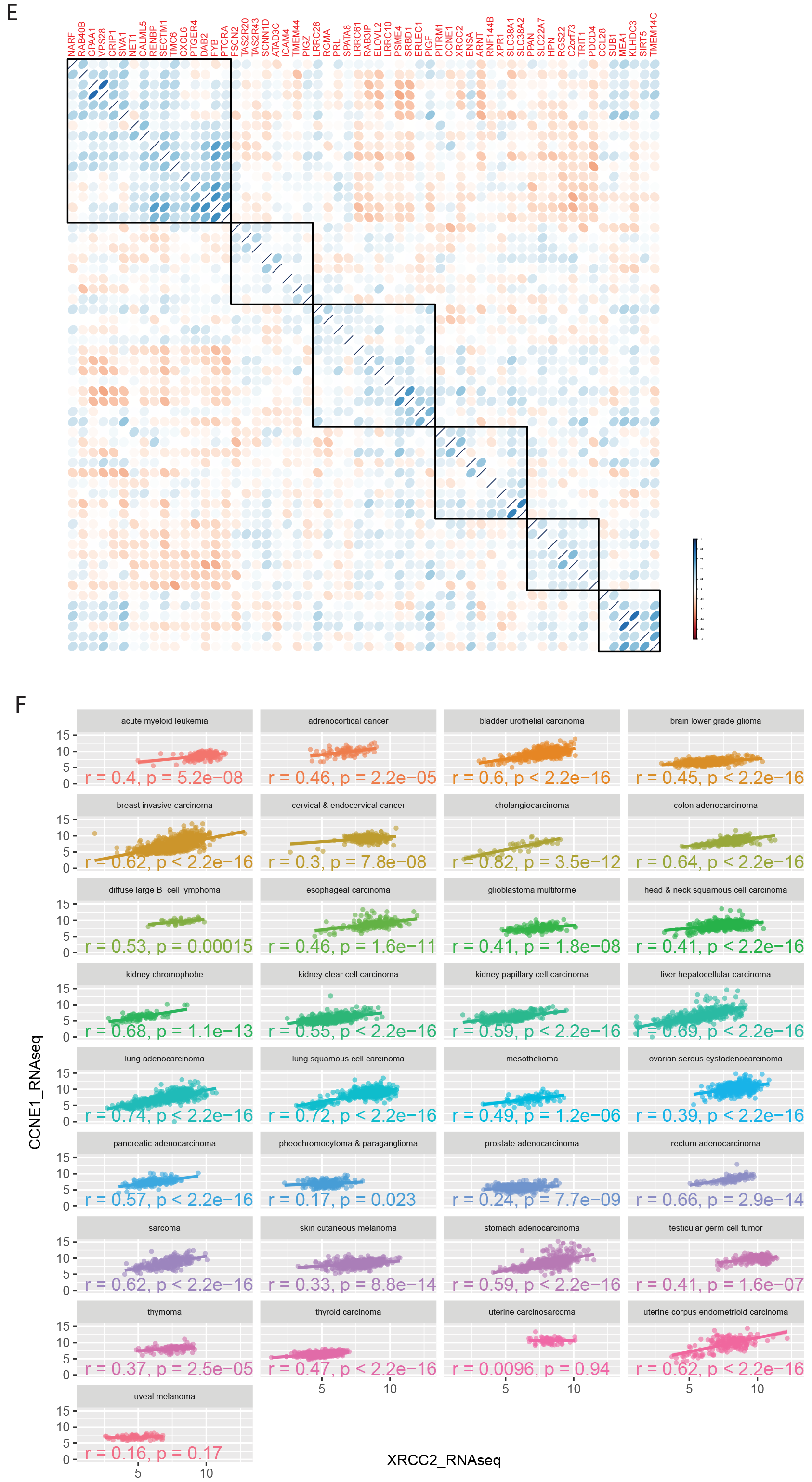

**Figure S2.**
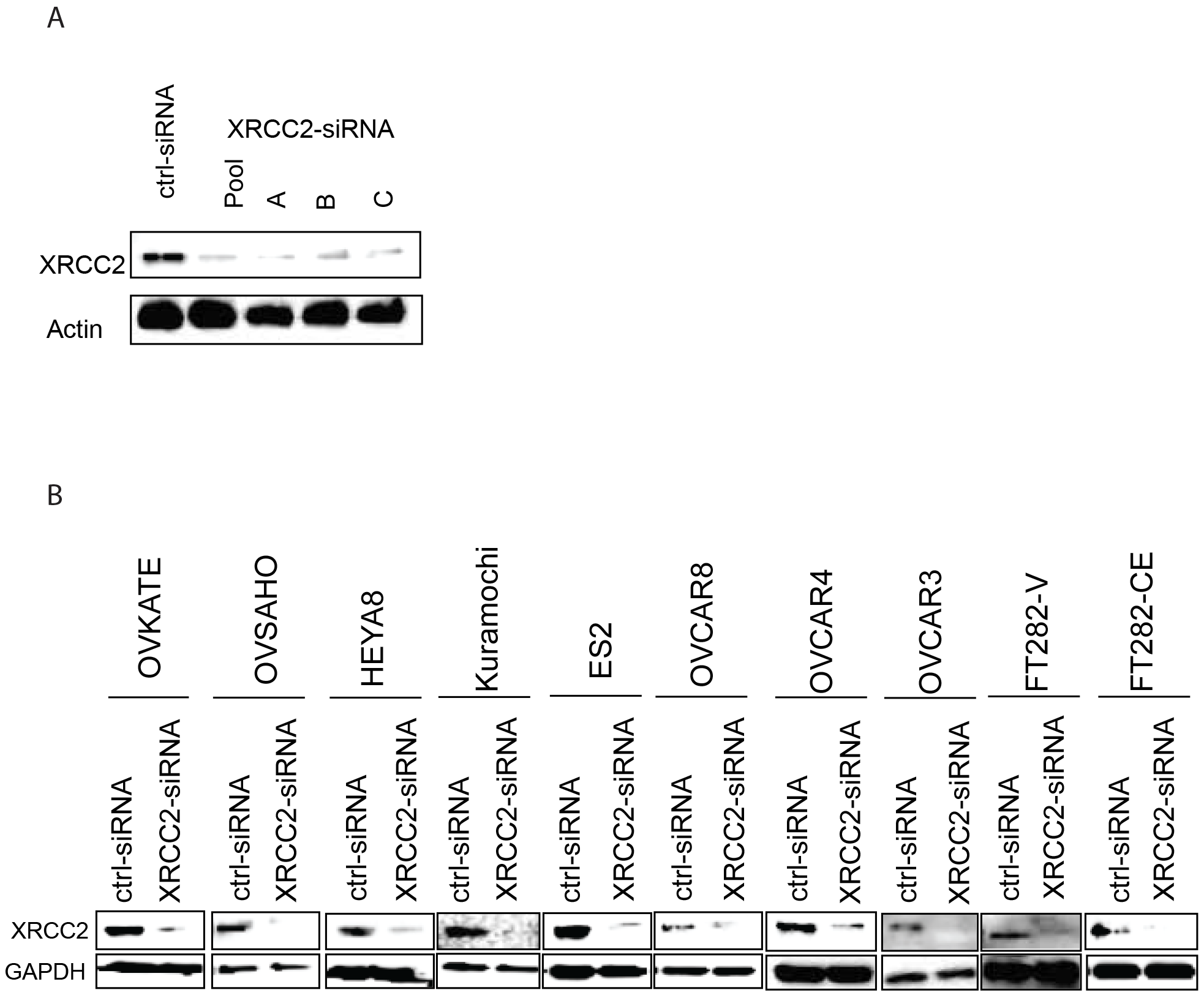
A) Western blot analysis of XRCC2 and beta-Actin in FT282-CE cells transfected with control siRNA, XRCC2 siRNA A, B, C or the pool of A,B and C. B) Western blot analysis of XRCC2 and beta-Actin in an ovarian cancer cell line panel after siRNA knock down of XRCC2. C) Violin plot of the XRCC2 dependency score (Demeter, RNAi) of the Achilles dataset comparing different cancer types. D) Scattered blot of XRCC2 dependency score (Demeter, RNAi) against CCNE1 RPPA (upper panel) and CCNE1 CNA level (lower panel). E) Left: dot plot of XRCC2 dependency score (Demeter, RNAi) of the Achilles dataset in cell lines that harbor high (CNA>0.45) against low CCNE1 copy number levels. right: dot plot of XRCC2 dependency score (Demeter, RNAi) of the Achilles dataset in cell lines that express high (RPPA>1) or low CCNE1 protein levels. F) Scatter plot of XRCC2 dependency score against CCNE1 CNA levels dataset separated in different cancer types. Shown is Pearson correlation for each cancer type. G) Waterfall plot of Pearson correlation between XRCC2 dependency score and CCNE1 CNA levels in each cancer type.

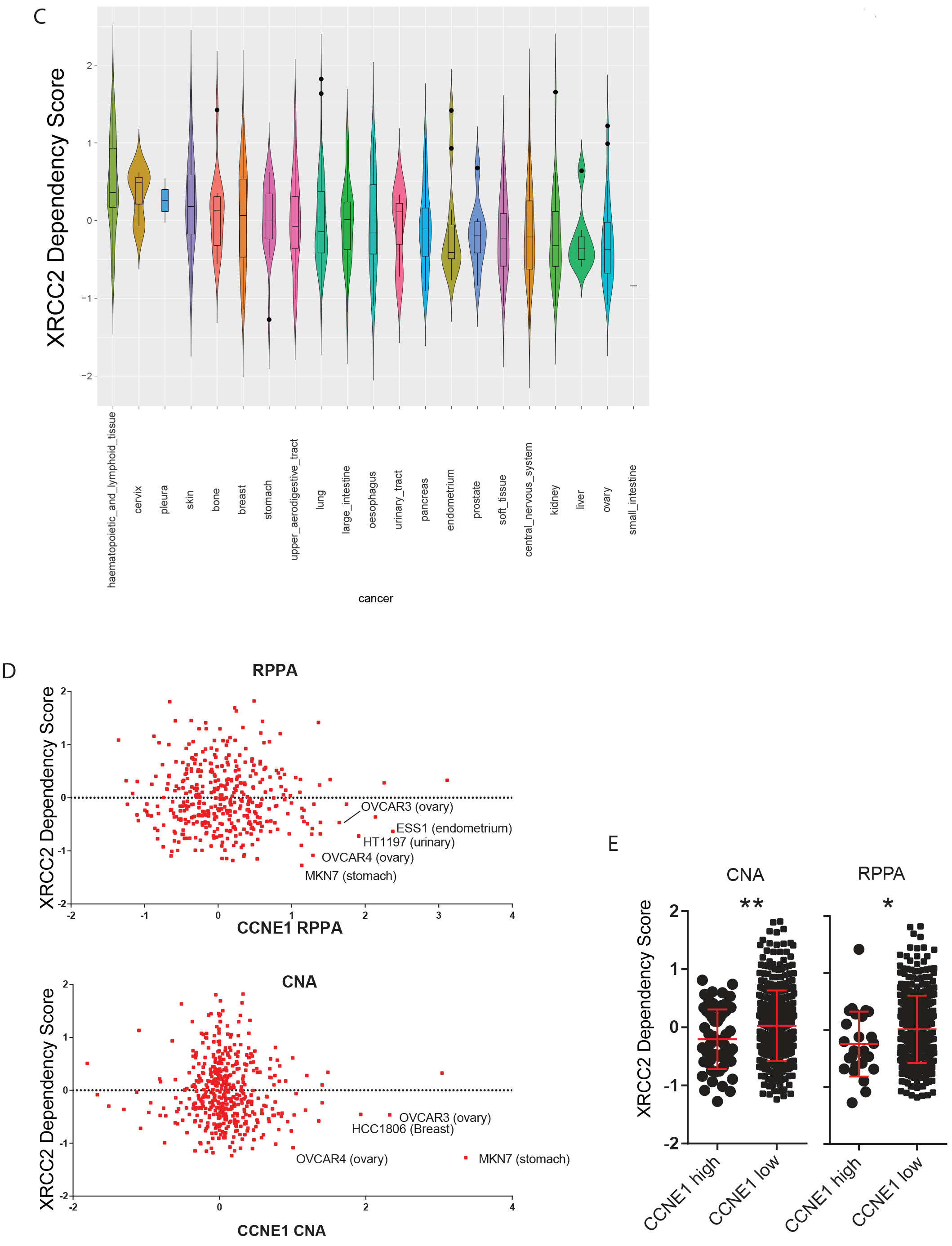

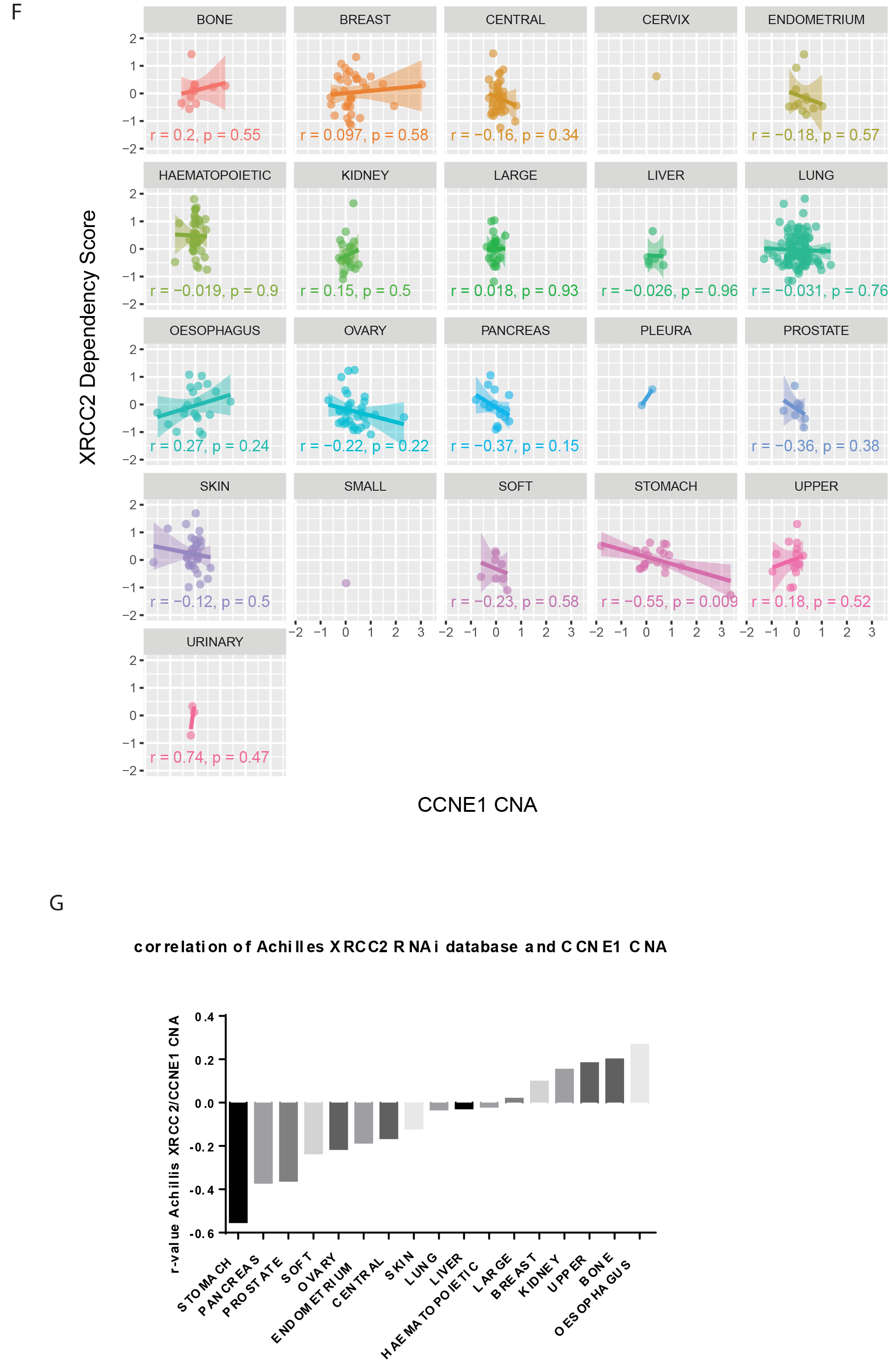

**Figure S3.**
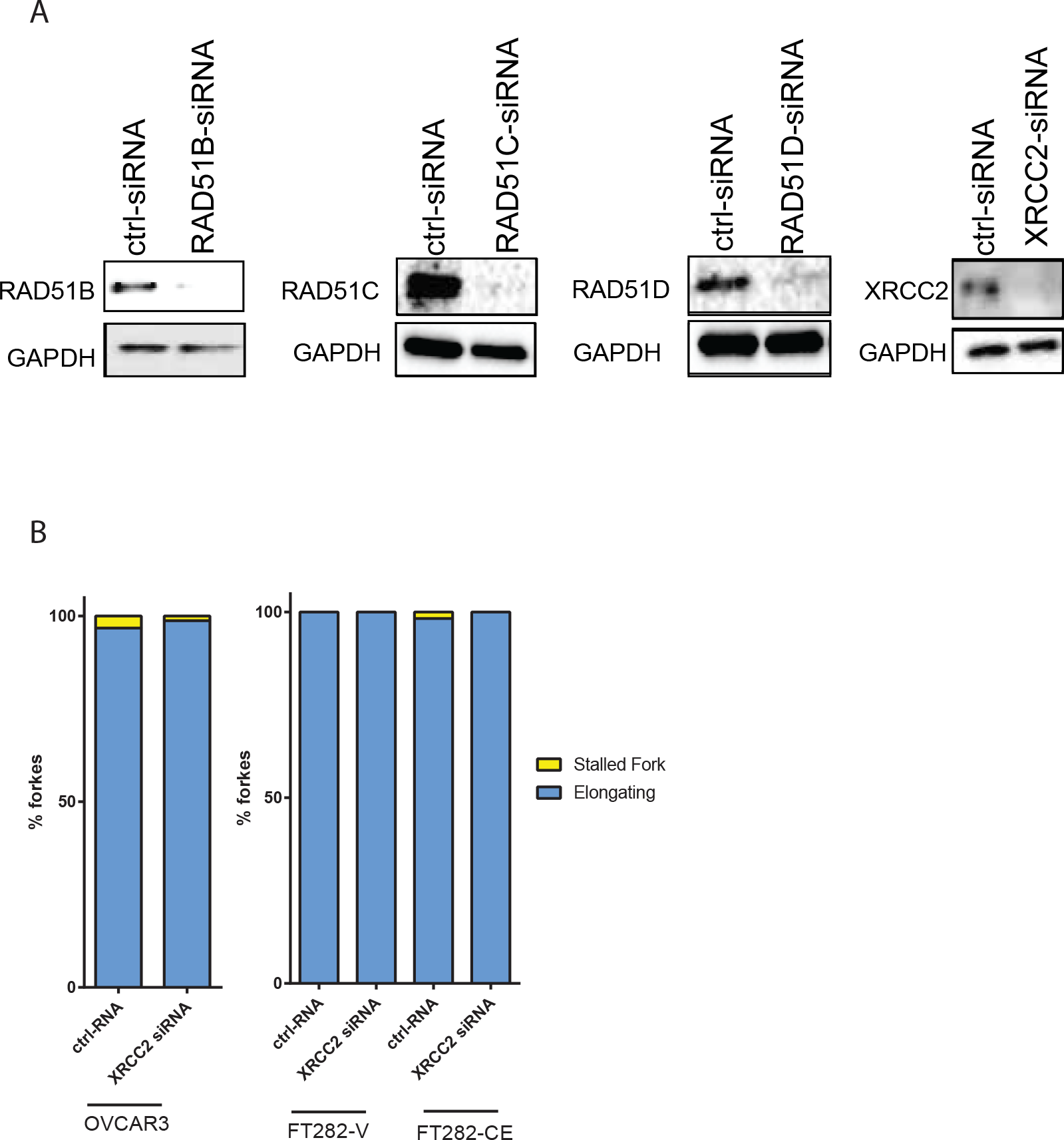
A) Western blot analysis of RAD51B, RAD51C, RAD51D, XRCC2 and beta-Actin in OVCAR3 cells cells transfected with control siRNA, RAD51B, RAD51C, RAD51D or XRCC2 siRNA. B) Analysis of fiber spread assay from figure 3C and 3D of OVCAR3, FT282-V and FT282-CE cells for Elongated and Stalled forks.

**Figure S4.**
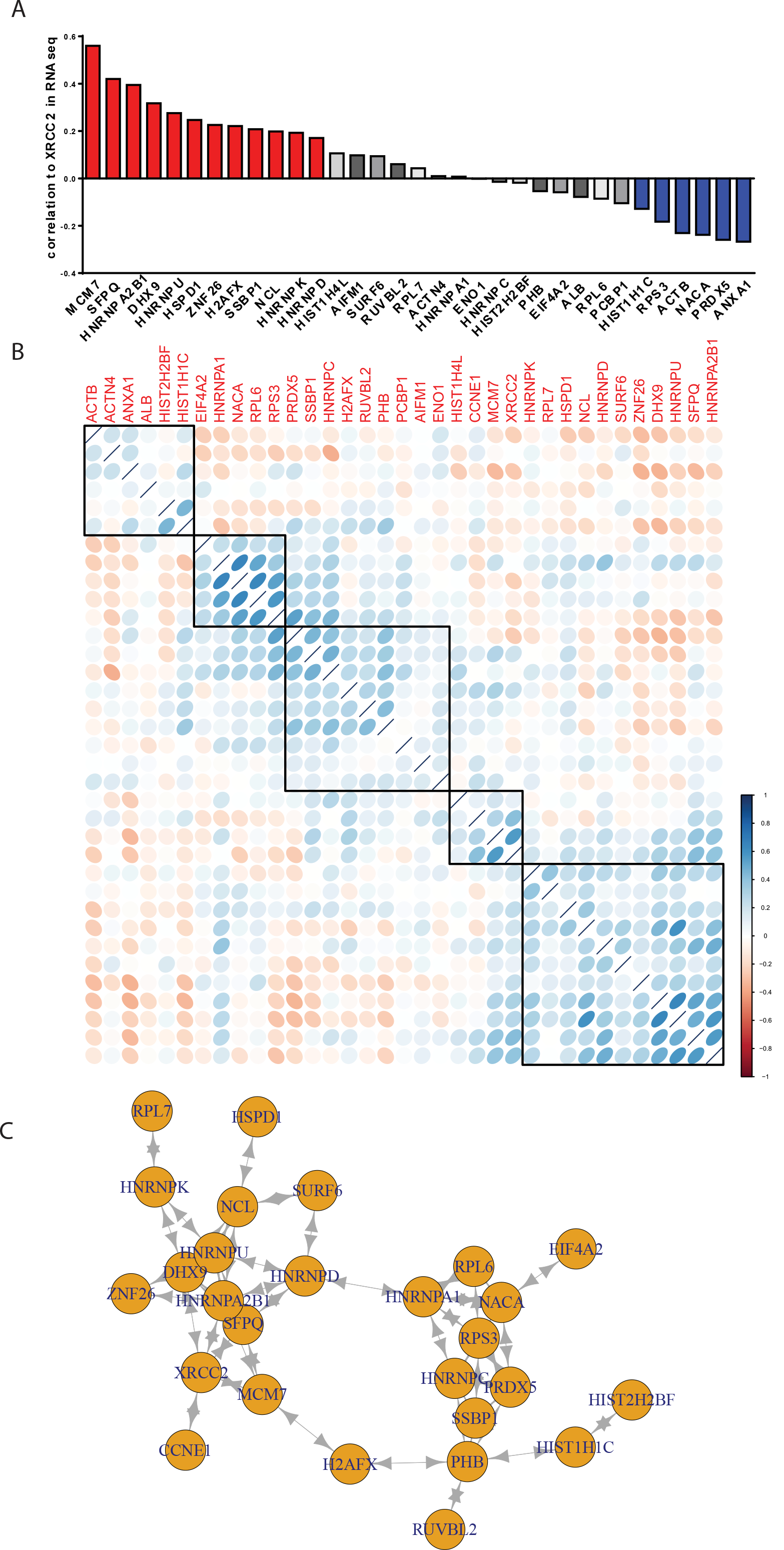
A) Samples from Figure 5A were assessed by mass spectrometry and identified genes were further analyzed for RNAseq expression in the TCGA ovarian cancer cohort and Pearson correlation to XRCC2 expression was analyzed. Red columns indicate a significant (p<0.05) positive correlation, whereas blue columns indicate a significant (p<0.05) negative correlation. B) Correlation matrix of the RNAseq expression of the identified genes and CCNE1 based on the TCGA ovarian cancer dataset and hierarchical clustered. C) Network analysis based on correlation matrix for genes that had a Pearson correlation higher than 0.3. D) Scattered blot of the RNAseq expression of *MCM7* against *XRCC2* expression in the TCGA Pan-cancer cohort separated for each individual cancer type. Depicted is pearson correlation. E) Correlation matrix of the RNAseq expression of MCM2, MCM3, MCM4, MCM5, MCM6, MCM7, XRCC2, RAD51B, RAD51C, RAD51D and CCNE1 based on the TCGA ovarian cancer dataset and hierarchical clustered.

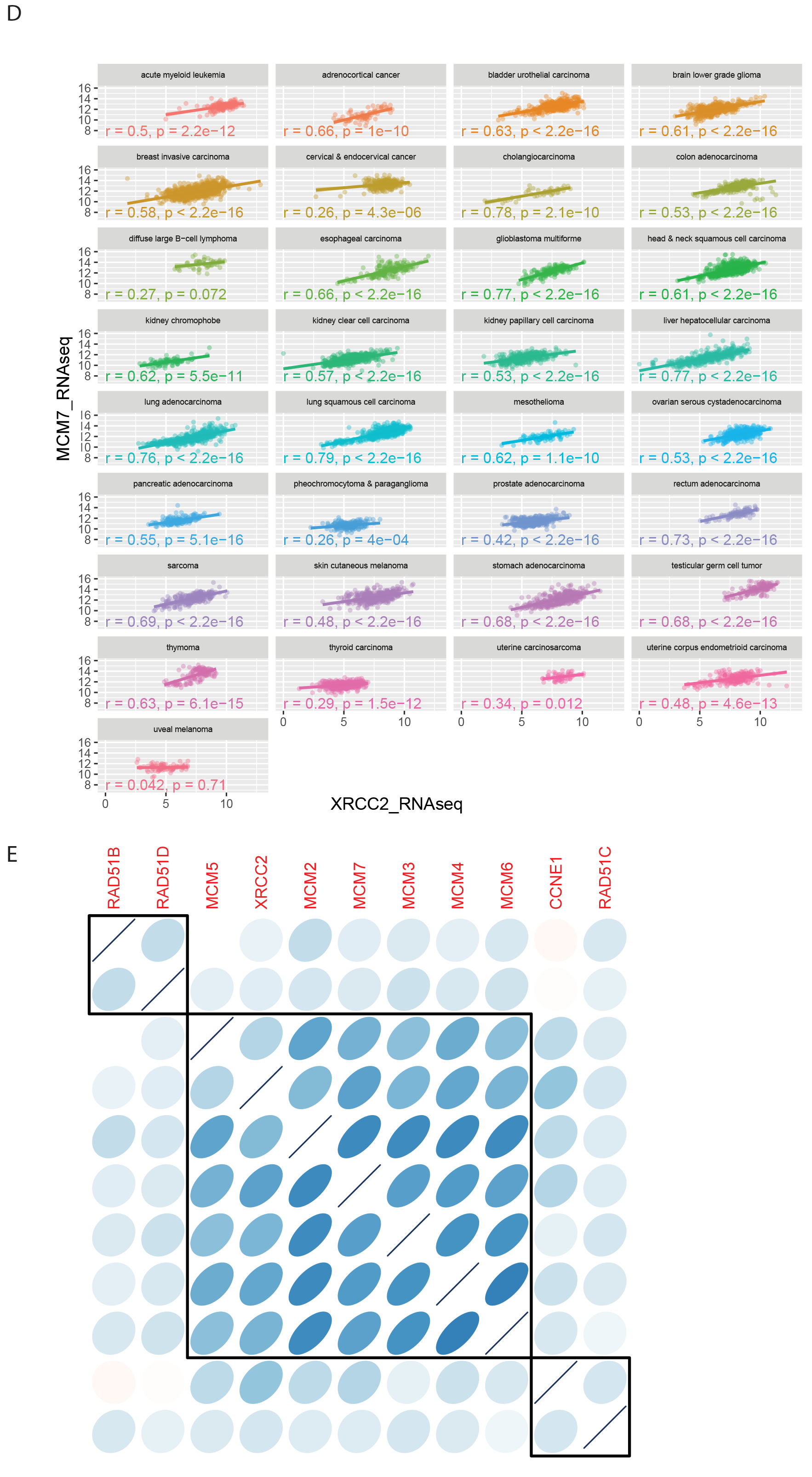

**Figure S5.**
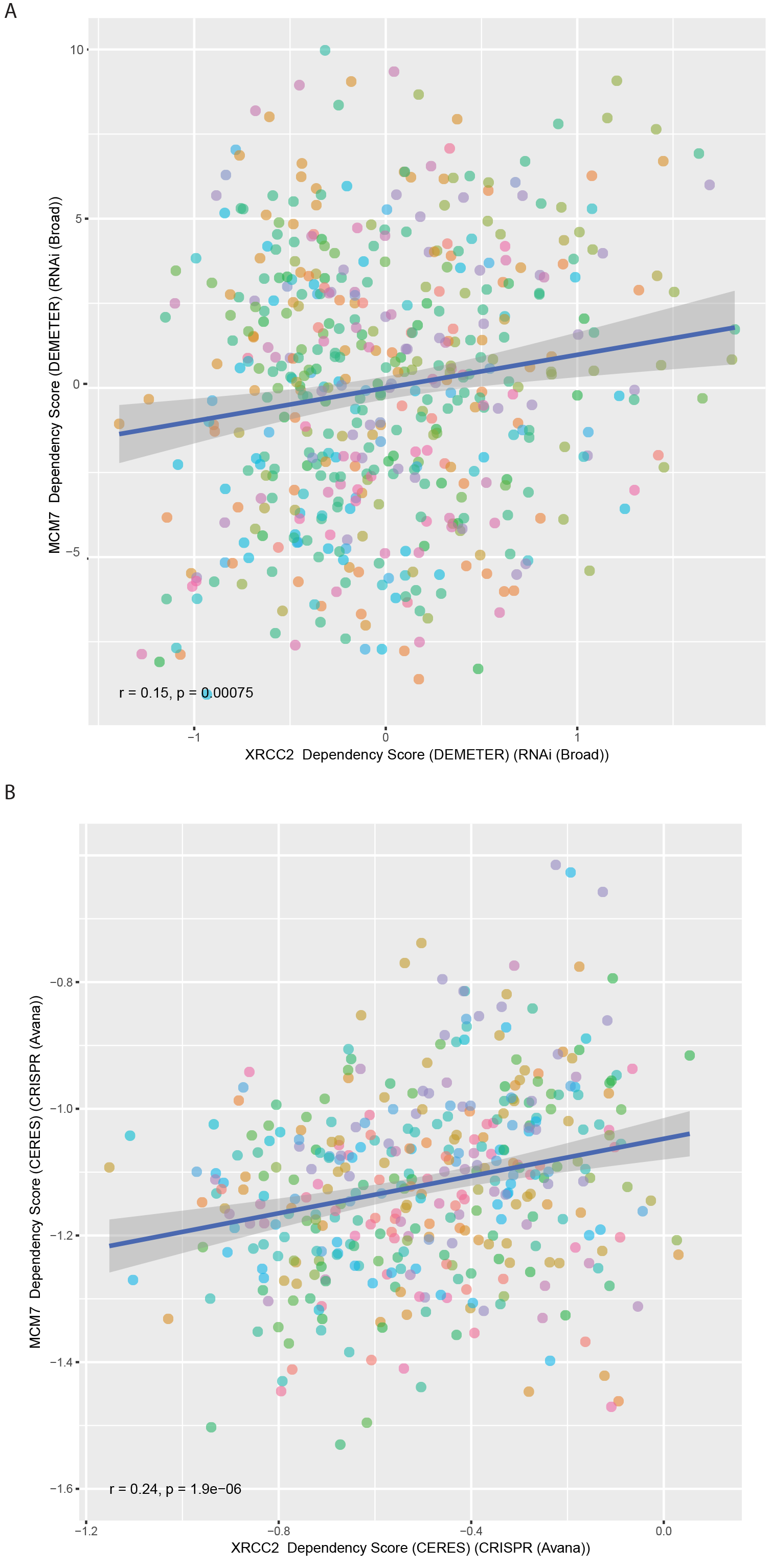
A&B) Scatter plot of MCM7 dependency score against the XRCC2 dependency score of the Achilles DEMETER(RNAi) (A, Pearson: r=0.15, p=0.00075) and CRISPR(CERES) (B, Pearson: r=0.24, p=1.9x10^-6^) dataset in all cell lines. C&D) Scatter plot of MCM7 dependency score against the XRCC2 dependency score of the Achilles DEMETER(RNAi) (C) and CRISPR(CERES) (D) dataset in separated in cancer types. E) Scatter plot of MCM7 dependency score against the XRCC2 dependency score of the Achilles CRISPR(CERES) dataset in ovary adenocarcinoma cell lines.

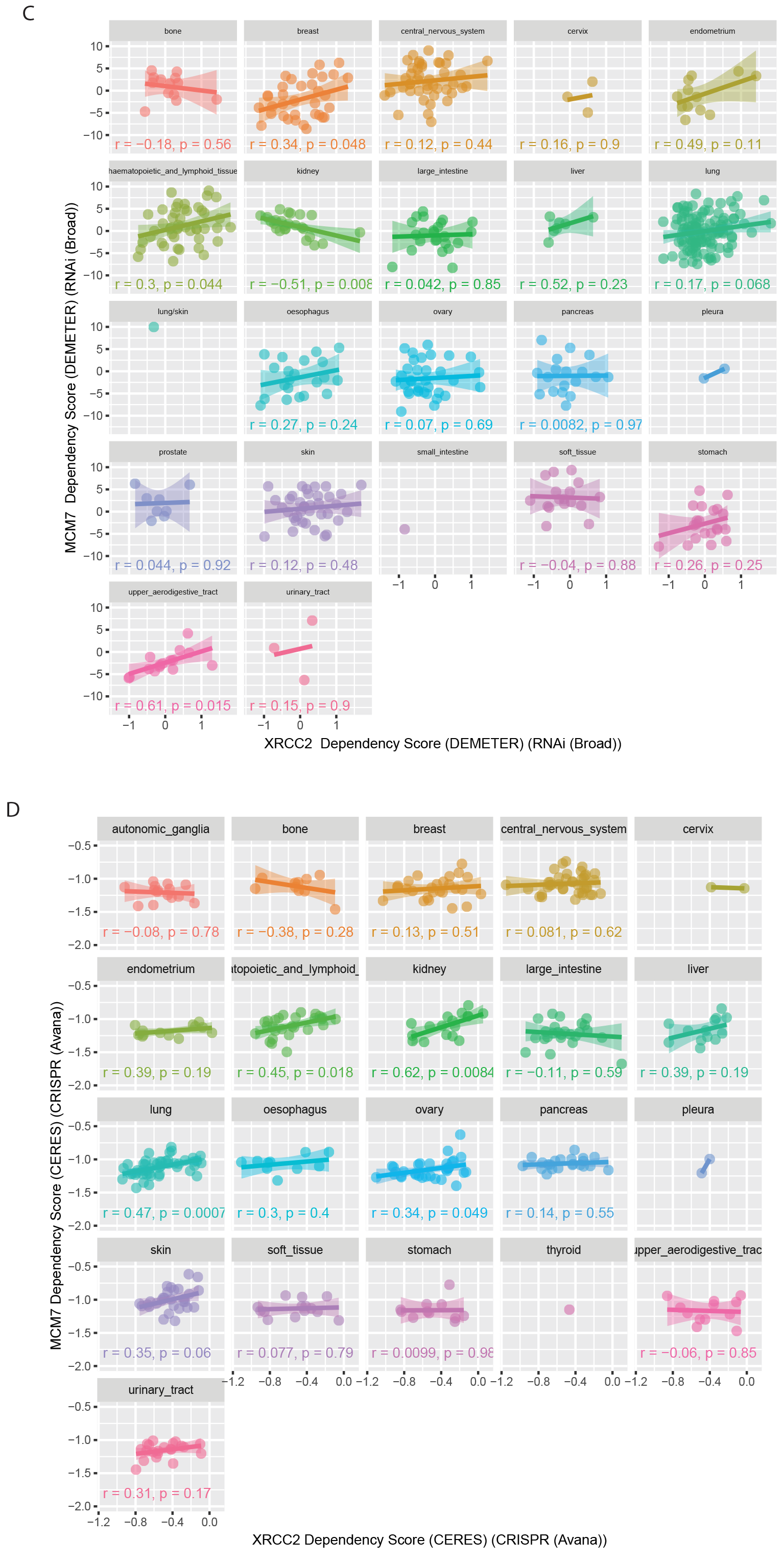

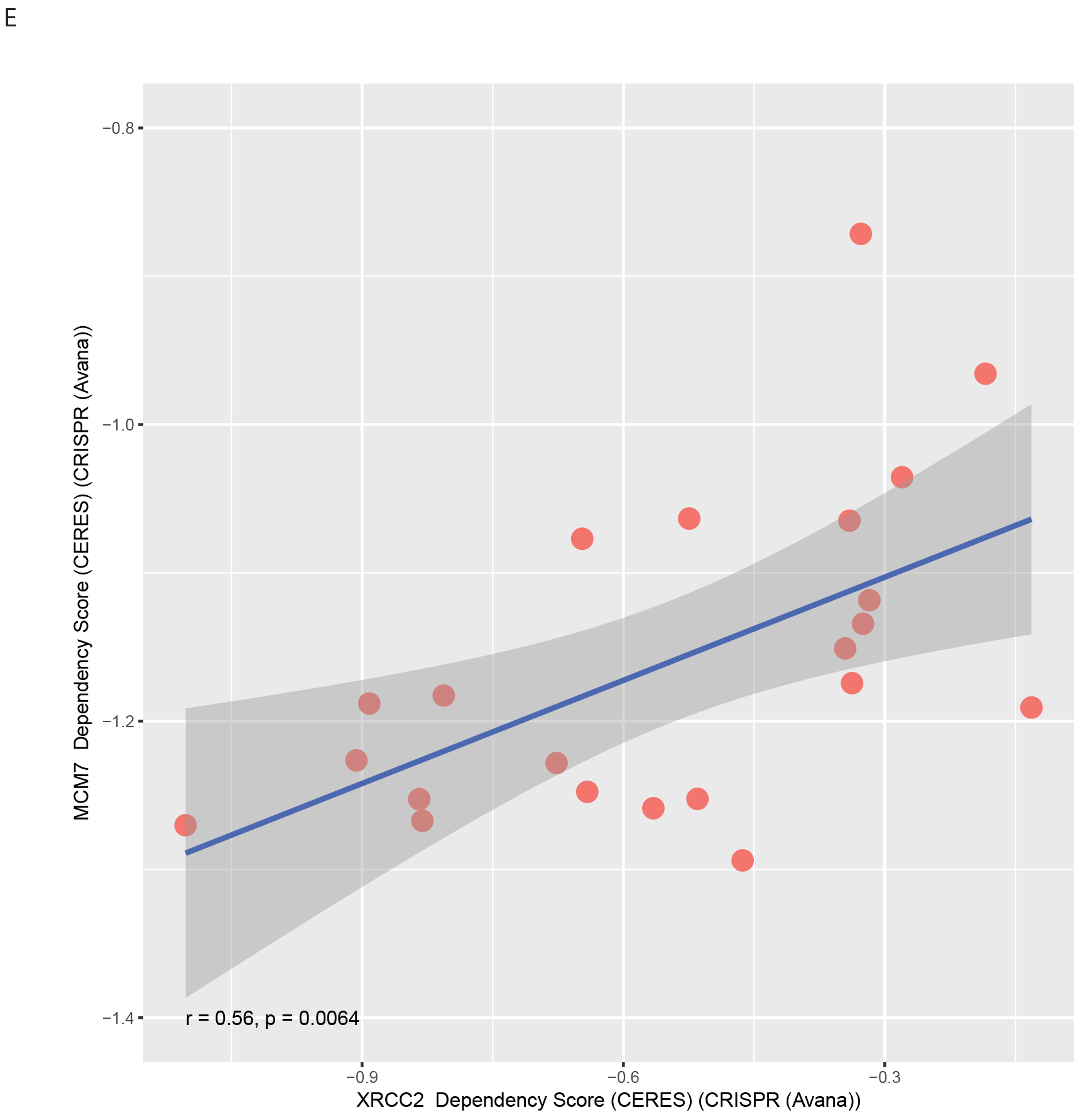

**Figure S6.**
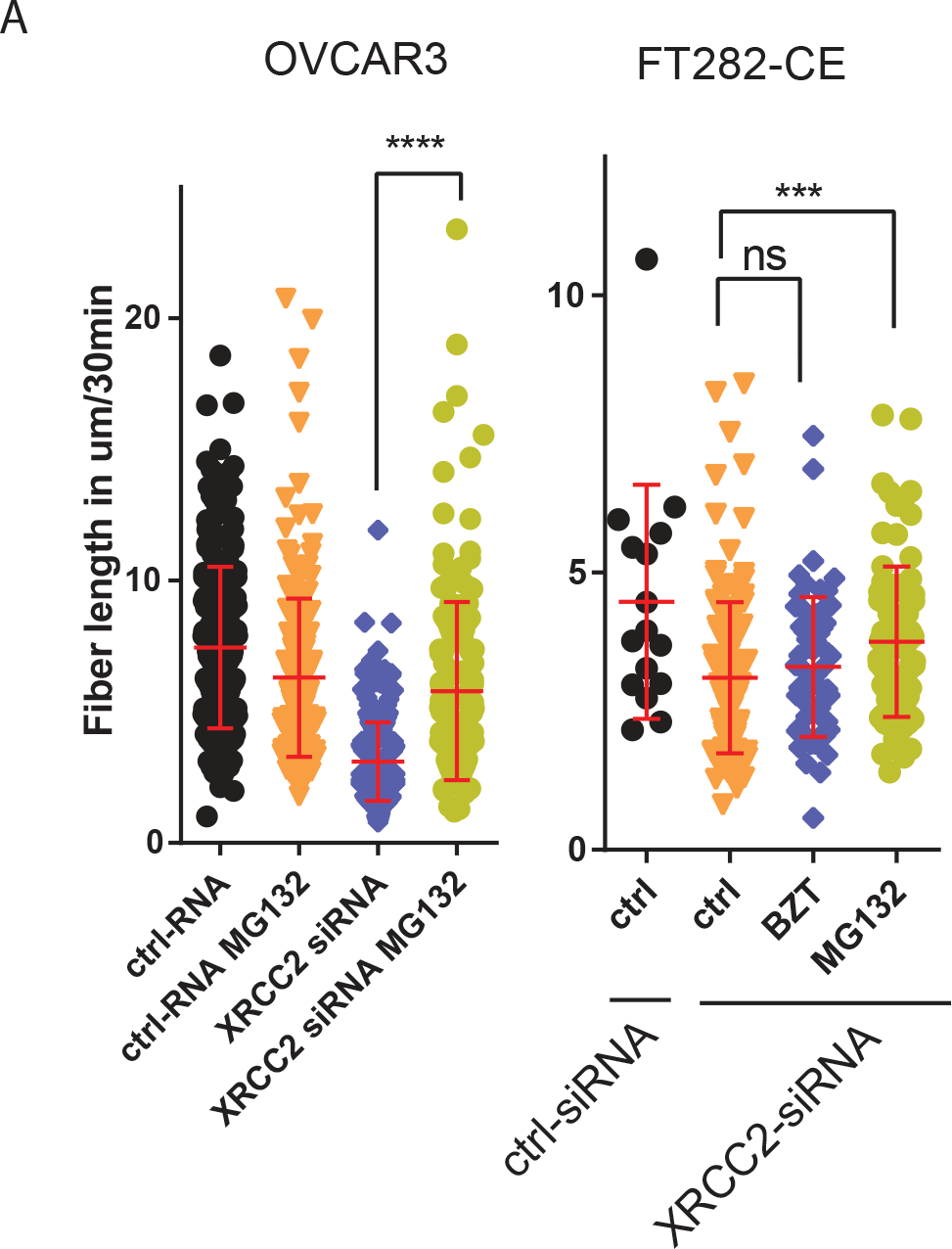
A) Fiber length analysis in OVCAR3 and FT282-CE measured after transfection of XRCC2. 16 hours before analysis, sampled were treated with proteasome inhibitors BZT or MG132. B&C) Scatter plot of VCP RNAseq expression against XRCC2 dependency score of the Achilles CRISPR(CERES) dataset in all cell lines (B: Pearson: r=-0.011, p=0.83) or separated in individual cancer types (C). D&E) Scatter plot of INT6 RNAseq expression against XRCC2 dependency score of the Achilles CRISPR(CERES) dataset in all cell lines (B: Pearson: r=-0.011, p=0.83) or separated in individual cancer types (C).

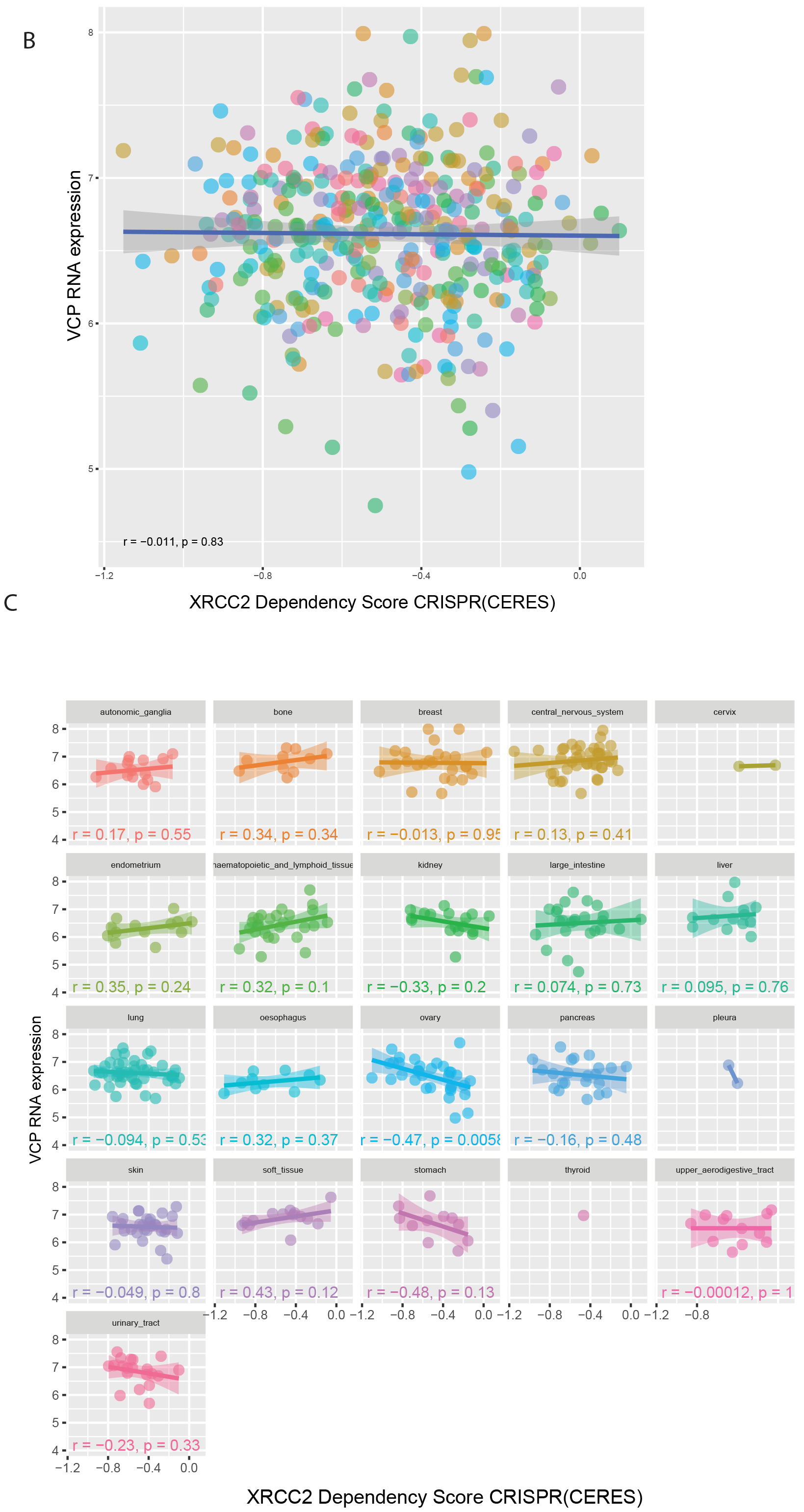

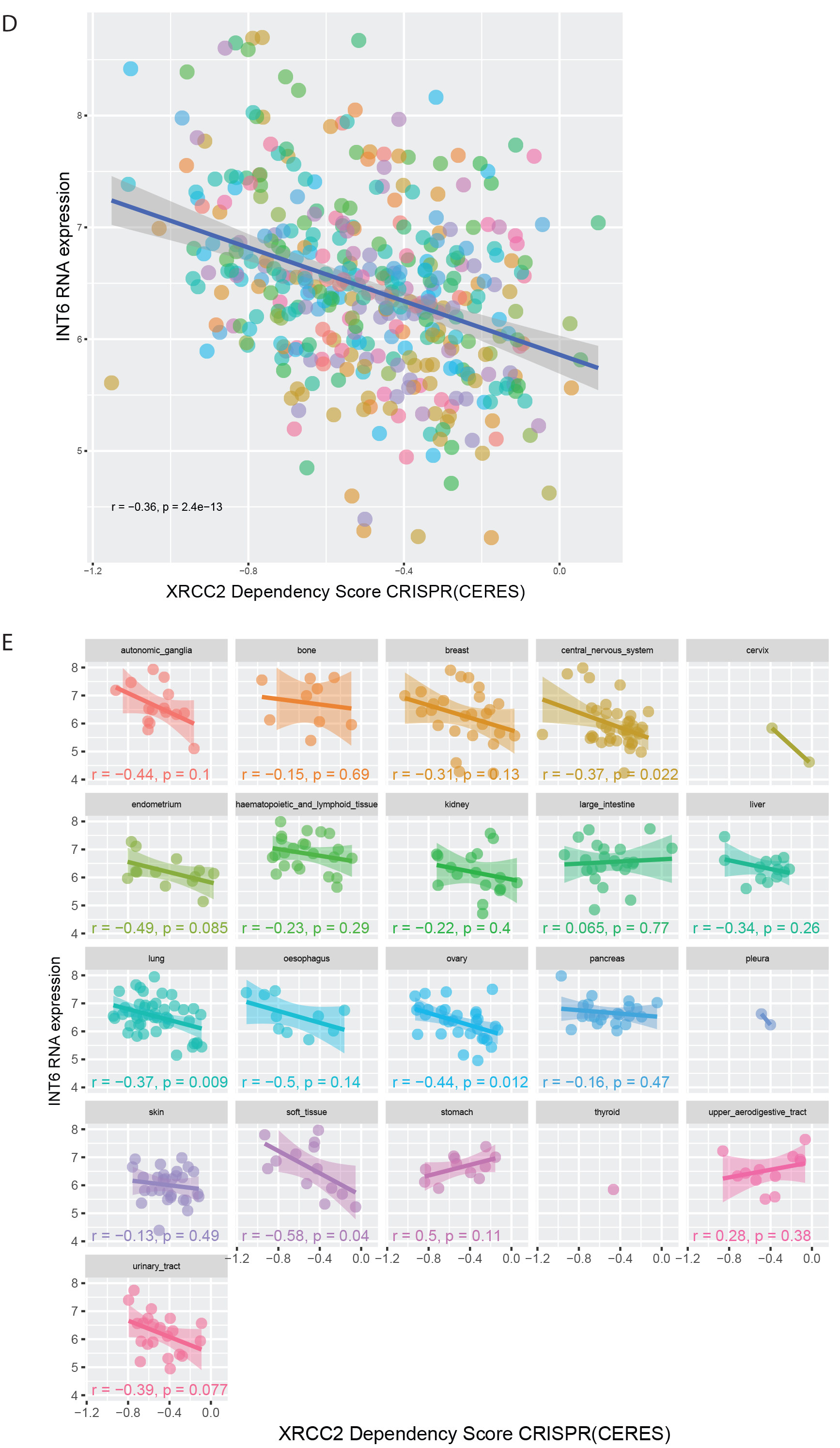

## STAR+METHODS

KEY RESOURCES TABLE

KEY RESOURCES TABLE
| Cell lines | cell line origins | Media |
| --- | --- | --- |
| FT282-V | Drapkin lab | DMEMF12+2%<br>USG |
| FT282-CE | Drapkin lab | DMEMF12+2%<br>USG |
| OVCAR3 | ATCC | RPMI<br>+10%FBS |
| OVCAR4 | NCI/DCTD | RPMI<br>+10%FBS |
| OVCAR8 | NCI/DCTD | RPMI<br>+10%FBS |
| COV318 | ATCC | RPMI<br>+10%FBS |
| OVSAHO | Gottfried Konecny | RPMI<br>+10%FBS |
| FUOV1 | DSMZ | RPMI<br>+10%FBS |
| Kuramochi | Gottfried Konecny | RPMI<br>+10%FBS |
| HEYA8 | ATCC | RPMI<br>+10%FBS |
| ES2 | ATCC | RPMI<br>+10%FBS |
| OVKATE | HSRRB | RPMI<br>+10%FBS |

| Antibody | company | lot number | species | methods/dilution | Antibody Registry |
| --- | --- | --- | --- | --- | --- |
| XRCC2 | Abcam | 180752 | R | WB 1:1000,<br>PLA 1:100 |  |
| XRCC2 | Sigma | SAB1402394 | M | IP 10µg/ml, IF<br>1:100 | AB_10645878 |
| Cyclin E1 | Cell Signaling | #4129 | M | WB 1:1000 | AB_2071200 |
| β-Actin | Cell Signaling | #3700 | M | WB 1:1000 | AB_2242334 |
| Phospho-H2A.X (Ser139) | Cell Signaling | #9718 | R | WB 1:1000 | AB_10121789 |
| RAD51B | Abcam | ab124675 | R | WB 1:1000 | AB_10973709 |
| RAD51C | Abcam | ab72063 | R | WB 1:1000 | AB_2177279 |
| RAD51D | Abcam | ab202063 | R | WB 1:1000 |  |
| FANCD2 | Abcam | ab108928 | R | WB 1:1000 | AB_10862535 |
| BRDU-Bio | Abcam | ab2284 | Shp | IP 20µg/ml | AB_302944 |
| BRDU/IdU | BD Bioscience | 555627 | M | Fibre 1:100 | AB_395993 |
| BRDU/CldU | Abcam | ab6326 | rat | Fibre 1:200, IF<br>1:100 | AB_305426 |
| RAD51 | Cell Signaling | #8875 | R | WB 1:1000 |  |
| POLD | Abcam | ab186407 | R | WB 1:1000 |  |
| MCM7 | Cell Signaling | #3735 | R | IP 10µg/ml, WB<br>1:1000, PLA<br>1:100 | AB_2142705 |
| MCM7 | Abcam | ab2360 | M | PLA 1:100 | AB_303010 |
| VCP | Cell Signaling | #2648 | R | WB 1:1000 |  |
| FK2 | Life Sensors | AB120 | M | PLA 1:100 |  |
| PCNA | Cell Signaling | #13110 | R | WB 1:1000 |  |
| MCM3 | Cell Signaling | #4012 | R | WB 1:1000 | AB_2235150 |
| MCM2 | Cell Signaling | #3619 | R | WB 1:1000 | AB_2142137 |
| control IgG | Cell Signaling | #5415 | M | IP 10µg/ml, |  |
| control IgG | Cell Signaling | #3900 | R | IP 10µg/ml, | AB_1550038 |
| GAPDH | Cell Signaling | #5014 | R | WB 1:1000 | AB_2107301 |

| Chemicals,<br>plasmids<br>and<br>recombinant<br>protein | company | lot number |
| --- | --- | --- |
| XRCC2-GST | Abcam | N9162 |
| MCM7 | Origene | TP313809 |
| Bromo-<br>deoxyuridine<br>(BrdU) | MP | 2100171 |
| Chloro-<br>deoxyuridine<br>(CldU) | MP | 2105478 |
| Iodo-deoxyuridine<br>(IdU) | MP | 2100357 |
| Doxorubicin | Tocris | 2252 |
| Cisplatin | TEVA | 00703-5747-11 |
| Neocarzinostatin | SIGMA | N9162 |
| MG-132 | SIGMA | M7449 |
| Bortezomib | SIGMA | 5.04314.0001 |
| Hydroxyurea | MP | 2102023 |
| GST | SIGMA | SRP5348 |
| PR619 | Life Sensor | SI9619 |
| 1,10-phenanthroline | Life Sensor | SI9649 |
| siRNA | company | lot number |
| XRCC2 | Origene | SR322227 |
| RAD51B | Origene | SR303968 |
| RAD51C | Origene | SR303967 |
| RAD51D | Origene | SR303970 |
| MCM7 | Origene | SR302840 |
| Plasmids | company |  |
| MCM7 | Vectorbilder |  |
| XRCC2-GFP<br>plasmid | Vectorbilder |  |
| Commercial<br>Assays | company | lot number |
| PLA | Sigma | DUO92101 |
| MTS-Assay | Promega | G3582 |

### Contact information for sharing

Further information and requests for resources and reagents should be directed to and will be fulfilled by the Lead Contact, Ronny Drapkin.

### Experimental procedures

#### Cell lines

The establishment of the FT282-V and FT282-CE cell line has been previously described (Karst et al, Cancer Research) and cultured in Dulbecco’s Modified Eagle’s Medium (DMEM)/Ham’s F-12 1:1 (Cellgro) supplemented with 2% Ultroser G serum substitute (Pall Life Sciences, Ann Arbor, MI, USA) and 1% penicillin/streptomycin. All cancer cell lines were cultured in RPMI 1640 (Invitrogen, Carlsbad, CA) supplemented with 10% fetal bovine serum (FBS, Atlanta Biologicals) and 1% penicillin/streptomycin (Invitrogen). All cells were grown inside an incubator maintaining 37°C and a 5% CO_2_-containing atmosphere.

#### Gain-of-function screen

The production of the lentiviral particles for the 588 ORF library has been previously described (Dunn et al., 2014). The 588 ORFs represent genes amplified in ovarian cancers as determined by The Cancer Genome Atlas (TCGA) dataset (REF)The positive and negative controls were obtained from the CCSB/Broad Institute ORF collection (Dunn et al., 2014). FT282-CE cells were plated at a density of 6,800 cells per well in 96-well plates and infected with-lentivirus generated from the lentiviral ORF plasmids. After 2 days, cells were transferred to 96-well ultra low attachment (ULA) plates and continuously inspected.

#### Immunoblotting and immunoprecipitation

Cells were lysed in RIPA buffer (25mM Tris-HCl pH 7-8, 150 mM NaCl, 0.1% SDS, 0.5% sodium deoxycholate, 1% Triton X-100 and protease inhibitors) for 3 h on ice and sonicated-for 3 ⨯ 30s at 20%. Protein content was quantified by BCA assay using the protocol enclosed in the Pierce BCA kit (#23228). 30μg were analyzed through a 4-15% gradient SDS-PAGE before transferred with the TurboBlot system (Bio-Rad) to PVDF membrane with a pore size of 0.2 μg. The membrane was blocked with 5% nonfat milk in TBST for one hour at room temperature. Blots were cut vertically, then incubated in primary antibody (diluted in Blocking buffer) overnight at 4°C. All antibodies and dilutions are listed in Supplementary Table S1. After washing, membrane was incubated in either anti-mouse or anti-rabbit HRP-conjugated secondary antibody (Cell Signaling, 1:1000) diluted in TBST for 1 hour. Proteins were detected using Clarity Chemiluminescent HRP Antibody Detection Reagent (Bio-Rad, #1705061) and visualized with a ChemiDoc imaging system (Bio-Rad).

#### TCGA dataset analysis

Data from the TCGA database were extracted and downloaded from the XENA portal of the University of California, Santa Cruz (http://xena.ucsc.edu/). The extracted copy number and RNAseq data from the TCGA ovarian cancer cohort, TCGA pan-cancer cohort and the Cancer Cell Line Encyclopedia were analyzed with the GraphPad Prism software and the R programming language.

#### Achilles dataset analysis

Data from the **Achilles** database were extracted and downloaded from the DepMap project at the Broad Institute (https://depmap.org/portal/). The data were analyzed with the GraphPad Prism software and the R programming language.

#### R programming language

For scatter plots we used the packages ggplot2, reshape2, ggpmisc, ggpubr. For correlation matrix we used the package corrplot and for the network analysis we used the packages igraph and MASS.

#### Viability assay

Cell viability was monitored after 72 hours using a CellTiter 96^®^ Non-Radioactive Cell Proliferation Assay (Promega, Mannheim, Germany). Each assay was performed in triplicates and repeated at least 3 times. Data are presented by means ± SD. Statistical and significant differences were determined by ANOVA with post-hoc analysis. Cells were additionally stained with crystal violet to count the remaining attached cells.

#### Short-interfering RNA transfection

A pool of three short-interfering RNA (siRNA) duplexes of the trilencer-27 siRNA (Origene) was used to downregulate the corresponding protein expression.

As a negative control an unspecific scrambled trilencer-27 siRNA was used. Twenty-four hours before transfection, 1 × 10^5^ cells were seeded in six-well plates (except OVCAR3, where 2 × 10^5^ cells were used). Transfection of siRNA was carried out using Lipofectamin RNAiMax (Invitrogen) together with 10 nM siRNA duplex per the instructions. To transfect 10 cm dishes we multiplied our protocol with the factor five and to transfect one well of a 96 well plate we divided our protocol by 20.

#### DNA fiber length assay

1 × 10^5^ cells were plated in each of six wells one day before siRNA transfection. 48 hours after transfection, cells were labeled with CIdU (38 μM) for 30 minutes before washing and subsequent addition of either medium containing IdU (250 μM) for 30 minutes or 5mM HU for 120min or 24 hour. Cells were harvested, washed and re-suspended in 20-40μl cold PBS .2.5 jl of the re-suspended cells were directly mixed with 7.5 μl lysis buffer (200 mM TrisHCl pH7.4, 50 mM EDTA, 0.5% SDS) on a slide. Slides were tilted manually 30°- 45° to allow the drop to run slowly down the slide. Once dried, the DNA was fixed in cold (−20C) methanol/acetic acid (3:1) for 5 minutes. Slides were dried and rehydrated 2⨯5 minutes in PBS before the DNA was denatured in 2.5M HCl for one hour. Slides were washed 5⨯3 minutes in PBS, blocked for 30 min (2% BSA 0.1% Tween 20 in PBS) and incubated with the primary antibodies mouse anti-BrdU/IdU (1:100, BD Bioscience) and rat anti-BRDU/CIdU (1:200, Abcam) for 2.5 hours at room temperature. Samples were washed 5x3 minutes in PBS containing 0.2% Tween-20 and one time briefly in blocking solution before incubated with secondary fluorescence antibodies (antimouse alexa 488, 1:300, Molecular Probes, anti-rat Cy3, 1:300 Jackson Immuno Research) diluted in blocking solution. After one hour, slides were washed 5x3 minutes and air dried in the dark before mounted with coverslip and 20 μl Antifade Gold (Invitorgen). Fibers were examined using a fluorescence microscope (Olympus) with a 100⨯oil immersion objective and analyzed using ImageJ software.

#### BrdU immunoprecipitation at Replication Forks

To immunoprecipitate the on-going replication forks, cells grown on two 15cm dishes cells were labelled with 100 μM BrdU (Sigma) for 30 min. The co-immunoprecipitation was adapted from the publications Yu et al. and Petermann et al., (Petermann et al., 2010; Yu et al., 2014). Cells were washed once with PBS and crosslinked with 1% formaldehyde for 10 min at RT before glycine was added to a final concentration of 0.125 M for 5 min at RT. After washing cells two times cold PBS cells were scraped from the plate and spun down at 5,000 g for 5 minutes. Cytoplasmic soluble protein fraction was removed by incubating cells with the first buffer (10 mM HEPES [pH 7], 50 mM NaCL, 0.3 M sucrose, 0.5% (v/v) Triton X-100, and protease inhibitor cocktail) for 10 min on ice and then spun down with 5,000 g for 5 min. To remove the nuclear soluble proteins, the pellets were incubated with nuclear buffer (10 mM HEPES [pH 7], 200 mM NaCL, 1 mM EDTA, 0.5% NP-40, and protease inhibitor cocktail) for 10 min on ice and spin down at 5,000 g for 5 min. Pellets were washed twice with cold PBS before DNA was digested to allow anti-BRDU antibody binding with 100 U/ml DNaseI (NEB) at 37°C for 50 min. After washing twice with cold PBS, the pellets were resuspended in lysis buffer (10 mM HEPES [pH 7], 100 mM NaCL, 1 mM EDTA, 1% NP-40, and protease inhibitor cocktail), sonicated for 3 times 30s and spun down at 13,000 rpm for 5 min. Total 1000 μg protein was incubated with 2 μg of biotin labeled anti-BrdU antibody (Abcam) for 20 hrs at 4°C in the dark. Streptavidin magnetic beads (Thermofisher) were added for one hour before beads were washed twice with cold lysis buffer. The proteins were eluted with 50 μl 2⨯SDS sample buffer by incubating for 15 min at 95°C and analyzed by western blot analysis.

#### Immunofluorescence

For immunofluorescent analysis, cells were grown overnight in a 96-well Cell Imaging Plates (Eppendorf). The procedure was adapted from Yu et al. (Yu et al., 2014). Cells were washed once with PBS and then permeabilized with 0.5% Triton X-100 for 10 minutes, and washed two times with PBS to remove soluble proteins. Cells were then fixed in 4% (v/v) paraformaldehyde in PBS for 20 minutes at room temperature. Cells were blocked with super-block buffer (Thermo Scientific) and incubated with primary and then secondary antibodies at 37°C for 1 hour. The following primary antibodies were used: XRCC2 (Abcam and Sigma) and MCM7 (Cell Signaling Technology). Detection was performed using secondary antibodies conjugated to Cy3 and Alexa Fluor Dyes (Molecular Probes). Cells were then mounted with DAPI containing Flouromount-G (Sigma-Aldrich) prior to microscopy using a Nikon E400 microscope under x40 magnification. For co-localization analysis with BrdU-labeled replication forks, cells were cultured in media with 100 μM CldU for 30 min, then immediately washed twice with PBS and processed for staining. The immunofluorescence was performed as described above with the exception that the primary antibodies against XRCC2 and BrdU were incubated in a buffer containing DNase I (0.5% BSA, 0.5 × PBS, 30 mM Tris-HCL pH7.5, 0.3 mM MgCl2, 100 U/ml DNase I) at 37°C for 50 min.

#### PLA MCM7-XRCC2 and MCM7-FK2

For the proximity ligation assay (PLA) analysis, cells were grown overnight on in 96-well Cell Imaging Plates (Eppendorf). Cells were fixed in 4% paraformaldehyde in PBS for 20 minutes, washed in PBS and permeabilized with 0.5% Triton X-100 for 10 minutes. Cells were blocked with super-block buffer (Thermo Scientific) and incubated with primary and antibodies at 4°C for overnight as follows: XRCC2 (Abcam) and MCM7 (Abcam) or the anti-poly-ubiquitin antibody FK2 (Life Sensors) and MCM7 (Cell signaling). The assay was performed using the Duolink kit (Sigma-Aldrich) according to the manufacturer’s protocol-Background control was produced by performing parallel experiments in which the 2 primary antibodies were left out of the procedure. Cells were then mounted with DAPI containing Flouromount-G (Sigma-Aldrich) and imaged using a Nikon E400 microscope under ×40 magnification. Images were analyzed using ImageJ.

#### Co-immunoprecipitation

For immunoprecipitation of endogenous proteins, the chromatin fraction was isolated as described above for the BrdU immunoprecipitation, except that cells were not crosslinked and no DNAse was added. Immunoprecipitaion was performed with an anti-MCM7 (Cell-Signaling) or an anti-XRCC2 (Sigma) antibody over night at 4°C in the dark. Magnetic Dynabeads (Invitrogen, # 10001D) were added for 1 hour before washing with lysis buffer. The sample was eluted with 50 μl 2×SDS sample buffer by incubating for 15 min at 95°C followed by a Western blot. To analyze the XRCC2-GFP co-immunoprecipitation, nuclear extract was analyzed 72 hours after XRCC2-GFP transfection using the manufacturer’s protocol, followed by precipitation with magnetic GFP-trap beads for 20 hrs at 4°C in the dark (Chromotek). Beads were eluted with 50 μl 2×SDS sample buffer by incubating for 15 min at 95°C followed by a western blot or silver stain analysis.

#### Ubiquitin-pull down

For this assay we seeded three 10cm dishes with a confluency of 70-80%. One day after seeding the cells were transfected with XRCC2 or control siRNA for 24 hour before the protease inhibitors bortezomib (100nM) and MG-132 were added for 10 hour. Cells were harvested and lysed in RIPA buffer containing PR619 (LifeSensors Cat. No. SI9619), 1,10-phenanthroline (LifeSensors Cat. No. SI9649), phosphatase inhibitor and protease inhibitor. Cells were sonicated and centrifuged for 10 minutes at 13000 rpm. 1.5 mg protein lysate was then incubated with 20μl of TUBE2-Agarose resin (LifeSensors Cat. No. UM402) or control-Agarose resin and incubated for 4 hour at 4°C on a rotator. Afterwards, agarose was spun down at 5000 rpm for 5 minutes and the supernatant was collected as unbound fraction. Agarose beads were washed three times with TBST before beads were eluted with 50 μl 2×SDS sample buffer by incubating for 15 min at 95°C followed by a western blot analysis.

#### GST pulldown assay

For the GST pulldown assays, 0.03 nmol of either GST or GST-XRCC2 (Abcam) fusion protein was incubated for 2 h at 4°C with 0.03 nmol of full length recombinant MCM7 (Origene) in a final volume of 0.1 ml binding buffer (50 mM Tris-HCl pH 7.4, 150 mM NaCl, 10 mM MgCl_2_). After incubation, 0.01 ml of anti-glutathione-beads were added to each sample. The beads were washed three times and eluted with 2×SDS sample buffer by incubating for 25 min at 95°C and analyzed by Western blot.

#### Statistics

All correlation values were calculated with the Pearson correlation coefficient using linear regression. P-values were calculated with an unpaired, two-tailed t-test. *=p<0.05, **=p<0.01, ***=p<0.001, ****=p<0.0001

